# High-throughput enzymology reveals mutations throughout a phosphatase that decouple catalysis and transition state analog affinity

**DOI:** 10.1101/2022.11.07.515533

**Authors:** C.J. Markin, D.A. Mokhtari, S. Du, T. Doukov, F. Sunden, P.M. Fordyce, D. Herschlag

## Abstract

Using High-Throughput Microfluidic Enzyme Kinetics (HT-MEK), we measured over 9,000 inhibition curves detailing impacts of 1,004 single-site mutations throughout the Alkaline Phosphatase PafA on binding affinity for two transition state analogs (TSAs), vanadate and tungstate. As predicted by catalytic models invoking transition state complementary, mutations to active site and active site-contacting residues had highly similar impacts on catalysis and TSA binding. Unexpectedly, most mutations to more distal residues which reduced catalysis had little or no impact on TSA binding and many even increased affinity for tungstate. These disparate effects are accounted for by a model in which distal mutations alter the enzyme’s conformational landscape and increase occupancy of microstates that are catalytically less effective but better able to accommodate larger transition state analogs. In support of this model, glycine substitutions (rather than valine) were more likely to increase tungstate affinity, presumably due to increased conformational flexibility and increased occupancy of previously disfavored microstates. These results indicate that residues throughout an enzyme provide specificity for the transition state and discriminate against analogs that are larger only by tenths of an Ångström. Thus, engineering enzymes that rival the most powerful natural enzymes will likely require consideration not just of residues in and around the active site, but also of more distal residues that shape the enzyme’s conformational landscape and finetune the active site. In addition, the extensive functional communication between the active site and remote residues may provide interconnections needed for allostery and make allostery a highly evolvable trait.

**Significance Statement:** Transition state analogs (TSAs) resemble fleeting high-energy transition states and have been used to inhibit enzymes in nature and medicine, to learn about enzyme active site features, and to design and select new enzymes. While TSAs mimic transition states, they differ from actual TSs, and we exploit these differences here. Systematic TSA affinity measurements for 1,004 mutants of PafA (a model phosphatase enzyme) revealed effects in and around the active site that mirror their effects on catalysis, but TSA-binding and catalytic effects diverge more distally. These observations suggest that residues throughout an enzyme adjust its conformational landscape on the tenth-Ångström scale to optimize the active site for catalysis, rendering allostery more evolvable in nature but likely complicating enzyme design.

## Introduction

Transition state analogs (TSAs) have been used by nature (1, 2), in research to provide insights into the properties of enzyme active sites and enzymatic catalysis (3–6), and in medicine to provide highly effective drugs (6–10). Catalysis can be defined as preferential stabilization of a transition state over a reaction’s ground state (3, 11, 12). Thus, it is expected—and has been observed—that compounds with electrostatic or geometric features resembling the transition state but not the ground state bind more strongly to enzymes than substrates or standard inhibitors. For example, lactone and sugar analogs serve as TSAs of glycosidases by mimicking the planar geometry and positive charge accumulation at the reactive carbon in the oxycarbenium-like transition state (13, 14); for serine proteases, TSAs mimic the charge and geometry of their transition states by forming adducts with the nucleophilic serine (15). In addition, many strong inhibitors are bi-substrate analogs, mimicking the partial bond formed in the transition state and taking advantage of the lower entropic cost in binding a single transition-state-like species relative to the two substrates independently (16). Despite these many successes, it has also been widely recognized that no transition state analog is perfect—none can exactly replicate the geometry and electronic distribution of a species with partial bonds that exists at the peak of the reaction pathway (17).

Efforts to use TSAs to develop artificial enzymes highlight their strengths and weaknesses. The selection of antibodies with catalytic activity via TSA binding—so-called “catalytic antibodies”— demonstrated the fundamental concept of transition state complementarity of enzyme active sites (18, 19). Nevertheless, the catalytic power of antibodies falls well short of naturally-evolved enzymes (20, 21). These shortcomings have led to speculation about what may be missing in these proteins and their design. These observations, and the general inability to design de novo enzymes that rival the rate enhancements of the most proficient natural enzymes, further highlights possible gaps in understanding of catalytic features (22).

Research has understandably focused on the direct interactions that enzymes make with their substrates and transition states. Nevertheless, it is common to find residues remote from the active site that substantially influence enzyme activity. Despite considerable speculation about the roles of these remote residues, our knowledge of the mechanism by which they impact catalysis is limited, in part due to a dearth of data. We recently developed High-Throughput Microfluidic Enzyme Kinetics (HT-MEK), an approach that allows parallel expression, purification and quantitative kinetic and thermodynamic assays for multiple substrates, concentrations, and inhibitors for ~1500 enzyme variants in parallel. In prior work, we applied HT-MEK to study 1,036 mutants of a phosphomonoesterase of the alkaline phosphatase superfamily, PafA, revealing deleterious catalytic effects (on the chemical step) at 161 of its 526 residues, including many distal residues (23).

Here, we systematically interrogated the effects of these mutations throughout PafA on binding of the TSAs vanadate and tungstate (24, 25). By comparing TSA binding effects to effects on catalysis, we explore active site specificity. Our results reveal extensive specificity effects from many mutations remote from the active site, including mutations that are catalytically deleterious but enhance TSA binding. These results indicate that residues throughout the enzyme (and their interactions) are involved in precise positioning at the active site. Thus, engineering enzymes that rival the most proficient natural enzymes will likely require consideration not just of residues in and around the active site, but also residues throughout the protein that fine tune active site positioning.

## Results

PafA and other members of the Alkaline Phosphatase superfamily are extraordinary catalysts, providing rate enhancements of up to 10^27^-fold for the hydrolysis of phosphate monoester substrates, formally equivalent to a transition state stabilization of ~37 kcal/mol (24, 26). PafA is a 526-residue enzyme with a Rossmann fold core and an active site comprised of a bimetallo Zn^2+^ core, a nucleophilic threonine residue (T79), and three additional residues (K162, R164, and N100); these three residues and the amide backbone of T79 contact the phosphoryl oxygen atoms of the substrate (Fig. 1*A,B*). Following substrate binding, hydrolysis proceeds through nucleophilic attack by T79 to form a covalent phosphoryl enzyme intermediate (E-P) that is subsequently hydrolyzed by an attacking zinc-liganded hydroxide ion to generate inorganic phosphate (Pi) which is released from the enzyme (Fig. 1*A*). During both chemical steps, one Zn^2+^ ion within the bimetallo core activates the nucleophile and the other stabilizes the developing negative charge on the leaving group (Fig. 1*B*). Mutations of the substrate-contacting residues yield much larger deleterious effects on catalysis than on binding Pi (the reaction product and substrate for the reverse reaction), indicating that these side chains preferentially stabilize the transition state (24).

**Fig. 1.**
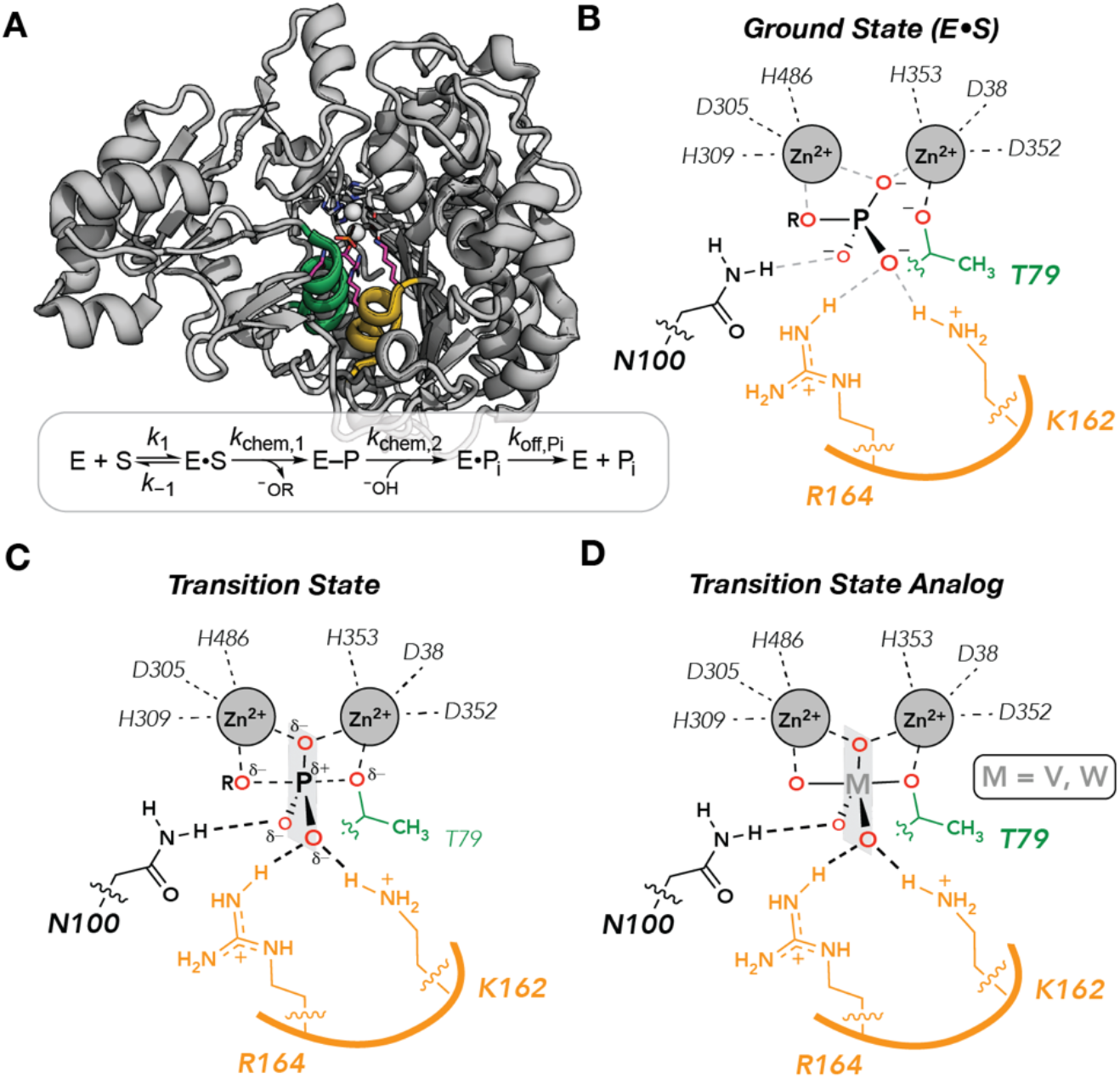
PafA catalysis and binding. **(A)** PafA active site and catalytic cycle showing helices containing the nucleophilic threonine (green) and active site residues K162 and R164 (orange). **(B,C,D)** Schematics of ground state **(B)**, transition state **(C)**, and transition state analog-bound state **(D)**. The semi-transparent plane contains the equatorial oxygens expected in the trigonal bipyramidal geometry of the transition state and of bound analogs.

Vanadate and tungstate bind strongly to many phosphoryl transfer enzymes and adopt trigonal bipyramidal geometries that mimic the reaction’s transition state (Fig. 1*C,D*) (25, 27–32). Consistent with expected TSA behavior, vanadate and tungstate affinity correlate closely with catalysis for 27 combinations of active site mutations in *E. coli* alkaline phosphatase and for four ablative active site mutations in PafA (24, 25). Nevertheless, crystal structures of these molecules in isolation and bound to proteins reveal differences in bond lengths and bond angles (see below). Here, we exploit this transition state mimicry and these subtle differences to learn more about the enzyme features that dictate molecular specificity in catalysis.

### HT-MEK reliably measures transition state analog binding affinities

To determine the impact of mutations throughout the enzyme on transition state analog binding affinity and specificity, we turned to HT-MEK (23). In HT-MEK, reaction chambers within valved microfluidic devices (Fig. 2*A*, left) are programmed with specific enzyme mutants by aligning devices to printed arrays of a library of plasmids encoding C-terminally eGFP-tagged PafA mutants. Following alignment, cell-free expression reagents are introduced into all chambers to express the mutants in parallel. After expression, eGFP-tagged mutants are recruited to anti-eGFP-patterned surfaces within each chamber to purify the enzymes in parallel (Fig. 2*A*, middle); measured eGFP fluorescence intensities along with an eGFP calibration curve report on the concentration of immobilized mutant enzyme. After immobilization, integrated valves that protect device surfaces make it possible to start and stop reactions as well as introduce fresh reagents without loss of surface-attached enzyme. To quantify TSA binding, we iteratively introduced the fluorogenic substrate 7-(dihydroxyphosphoryloxyl)coumarin-4-acetic acid (cMUP; (23)) in the presence of increasing TSA concentrations (7–13 concentrations ranging from 0.1 μM to 1 mM) and measured initial reaction rates via time-resolved fluorescence imaging (Fig. 2*A*, right). Increasing concentrations of vanadate and tungstate inhibited catalysis as expected, with behavior well fit by a competitive inhibition model (Fig. 2*B* and see below). Dissociation constants for vanadate, tungstate, and Pi obtained from these data for WT PafA and four active site mutants on-chip agreed well with values from traditional measurements (*r*^2^ = 0.95; Fig. 2*C*), as is also the case for kinetic constants (23, 24).

**Fig. 2.**
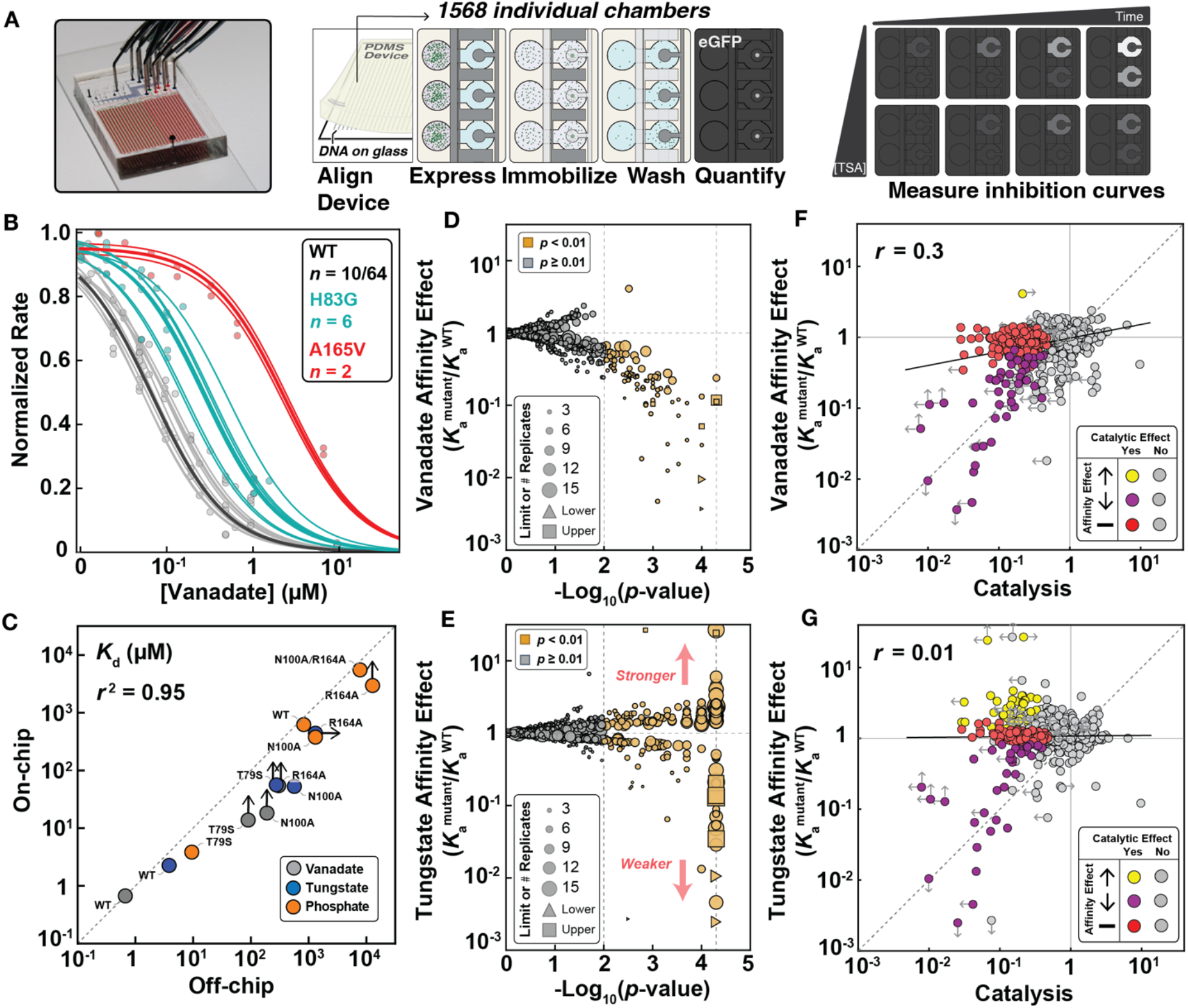
HT-MEK experimental pipeline and TSA binding measurements for PafA mutant library. (**A**) Device photo (left) and schematic showing experimental measurements of inhibition. (**B**) Representative vanadate inhibition curves. Thin lines indicate per-chamber curve fits; thick lines show fit behavior predicted by median *K*_i_ for WT (black), H83G (cyan), and A165V mutants (red); for WT, 10 of 64 total measurements are shown for clarity. (**C**) Comparison between dissociation constants (*K*_D_S) measured on- and off-chip for vanadate, tungstate, and Pi. Limits (denoted with arrows) were not used to calculate correlation coefficient. (**D**, **E**) Volcano plots of vanadate (**D**) and tungstate (**E**) association constants (*K*_a_^mutant^/*K*_a_^WT^) for PafA glycine and valine scanning libraries; marker size indicates number of replicates and limits are of affinity effects. (**F**) Comparison of vanadate affinity effects (*K*_a_^mutant^/*K*_a_^WT^) and catalytic effects (*k*_cat_/*K*_M,mutant_/*k*_cat_/*K*_M,WT_) for PafA mutants. (**G**) Comparison of tungstate affinity effects (*K*_a_^mutant^/*K*_a_^WT^) and catalytic effects. In (**F**) and (**G**), the points are colored by effect significance: mutants that do not differ significantly in catalysis from the WT construct are shown in gray; mutants with significant catalytic defects are colored based on vanadate (**F**) or tungstate affinity (**G**) effect (yellow, increase; purple, decrease; red, no effect). The dashed line is the identity line and the solid line the best-fit correlation line.

### Systematic HT-MEK measurements reveal hundreds of mutations that impact transition state analog binding

Over 17 experiments, we determined inhibition constants or limits for 1,004 of the 1,036 PafA variants in the presence of either vanadate or tungstate, corresponding to measurement of 1,944 equilibrium constants, with the remaining variants too catalytically impaired to quantify inhibition. We report inhibition curves for each mutant in each experiment in per-experiment and per-mutant reports in our Open Science Foundation data repository (https://osf.io/k8uer/), corresponding to a dataset of 9,278 inhibition curves. The high number of replicate measurements obtained via HT-MEK (*SI Appendix*,, Fig. S1) allowed us to resolve statistically significant affinity differences of <2-fold via bootstrap hypothesis testing (*p* < 0.01; Fig. 2*D*, *SI Appendix*, Fig. S2) (23, 33). Even with this high sensitivity, ~90% of the mutants showed wild-type-like vanadate inhibition (897/979 mutants), with 82 mutants decreasing affinity and one increasing affinity (Figs. 2*E*, *SI Appendix*, Fig. S2). Tungstate measurements revealed similar numbers of mutants with decreased affinity (76/964 mutants) (p < 0.01; Fig. 2*E*, *SI Appendix*, Fig. S3). Unexpectedly, a large number of mutants (155) significantly enhanced tungstate binding (Fig. 2*E*).

### Many mutations reduce catalysis without impacting transition state analog binding

Traditionally, a compound is considered a good TSA if changes to the enzyme (*e.g*. via mutation) or the analog (*e.g*. via chemical modification) yield equal magnitude changes in binding and catalysis (25, 30). Active site mutants of PafA and another Alkaline Phosphatase family member, *E. coli* alkaline phosphatase, give strong correlations between vanadate or tungstate binding and catalysis ((24, 25) & see below). Yet many PafA mutations (123) slowed the chemical step of catalysis without significantly weakening binding of either TSA: 50 of the 185 mutations that reduced catalysis weakened vanadate and/or tungstate affinity (Figs. 2*F,G*). This result is not a consequence of differences in statistical resolution between experiments; indeed, HT-MEK binding measurements have higher accuracy than HT-MEK kinetic measurements (due to errors in determining enzyme concentration that affect kinetic but not inhibition constants), so this differential is likely still larger (23). Furthermore, for mutations giving the largest catalytic effects (>10-fold), some yield similar reductions in vanadate and tungstate binding, but many yield little or no reduction in binding (Figs. 2*F,G*). Below, we evaluate which residues and substitutions give rise to equal or disparate effects and develop models to account for these results.

### Preferential impact on catalysis over vanadate binding from mutations distal to the active site

Prior mutations of active site residues measured off-chip gave nearly equal (within 2-3 fold) effects on catalysis and vanadate binding, consistent with expectations for removing interactions with a TSA that contribute to catalysis (Fig. 3*A*) (24). The sole exception is K162A, which gave a >10^6^-fold catalytic effect but a much smaller effect on vanadate binding (~10^2^). Moving out from the active site to residues that directly contact the catalytic residues or zinc ions (“second shell” residues), most mutants had very similar impacts on catalysis and affinity, with only four of the 39 measured preferentially impacting catalysis by 3-fold or more (D38G, G99V, Y306G, and N399V) (Fig. 3*B* and *SI Appendix*, Fig. S4). Second shell mutants include 12 catalytic Zn^2+^ ligand mutants; six of these mutants had measurable catalytic activity and only one, D38G, did not give similar deleterious effects on catalysis and vanadate binding (Fig. 3*B*, squares and *SI Appendix*, Table S1).

**Fig. 3.**
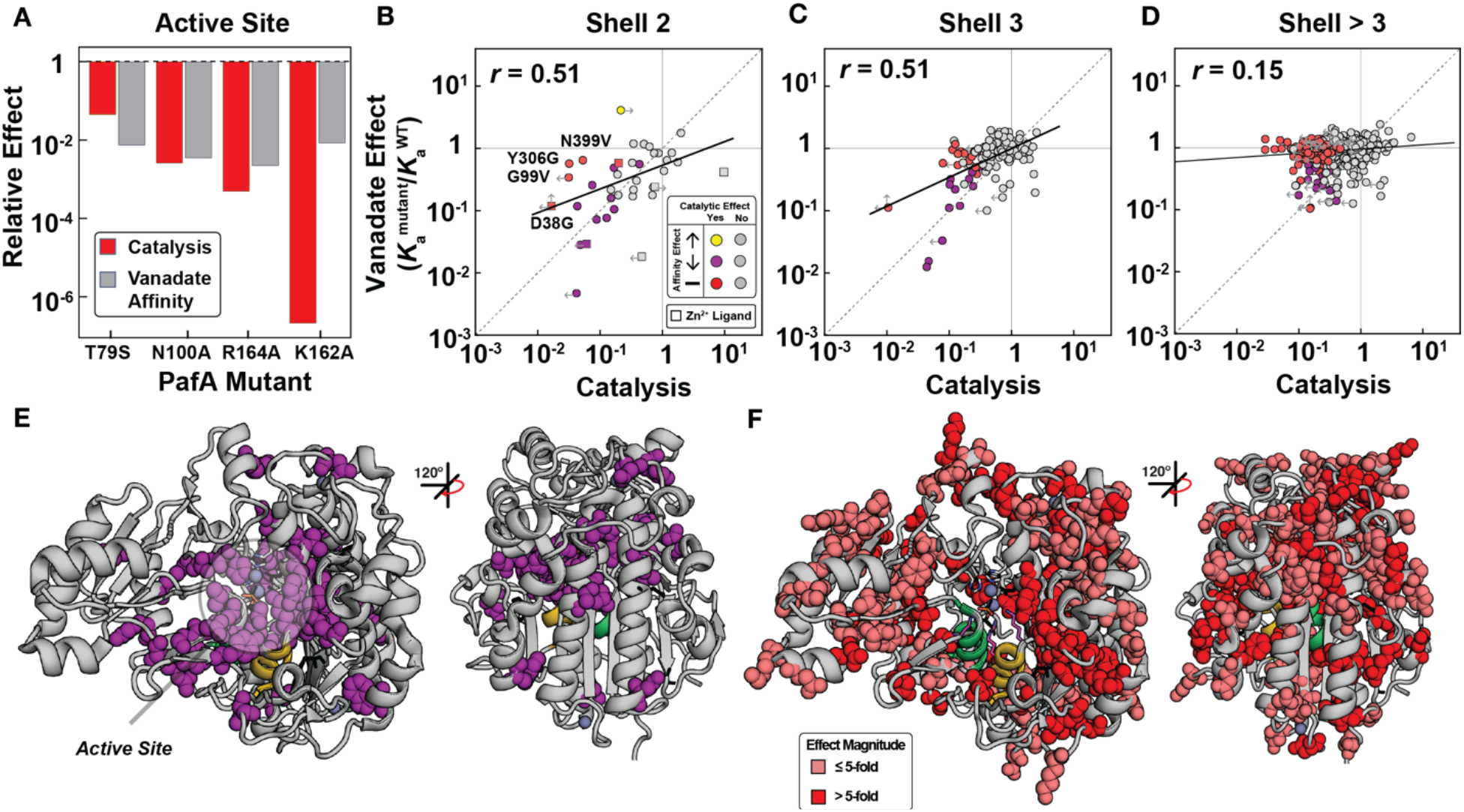
Effects of mutations on vanadate binding and catalysis as a function of distance from the PafA active site. (**A**) Comparison of catalytic and vanadate affinity effects for PafA active site mutants, measured previously (24). (**B-D**) Comparison of catalytic and vanadate affinity effects for mutants of residues in the (**B**) second shell, (**C**) third shell, and (**D**) more distal shells. Labeled points in (**B**) denote mutants with significant catalytic but not affinity effects. (**E**) PafA structure with spheres shown for residues with significantly deleterious catalytic and vanadate affinity effects when mutated to either glycine or valine (35 of 520 measurable residues) (**F**) PafA structure with spheres shown for residues residues with significant catalytic but not significant affinity effects (125 of 520 measurable residues). Catalytic data are from reference (23).

Moving further from the active site, we observed decreasing congruence between catalytic and vanadate-binding effects. In the third shell, few mutations gave similar effects (Fig. 3*C*, purple), with the majority affecting catalysis more than vanadate binding. Beyond the third shell (shells 4–8), there was essentially no correlation, even for distal mutants with large detrimental catalytic effects (Fig. 3*D* and *SI Appendix*, Fig. S5 and Tables S2,S3). This loss of congruence between catalytic and vanadate binding effects with increasing distance from the active site is also apparent from the decreasing slopes of correlation lines in Fig. 3*A–D* and decreasing correlation coefficients. Projecting these results onto the PafA structure provides a striking visualization of the diminishing correlation of catalytic and TSA-binding effects with distance from the active site (Fig. 3*E,F*).

### Increased tungstate binding from many distal mutations

We carried out the same analyses as above for vanadate with the second TSA, tungstate. Active site mutants again gave similar effects on catalysis and tungstate binding, but with enhanced agreement for the K162A mutant relative to that observed for vanadate (Fig. 4*A*) (24). Also as for vanadate, tungstate binding and catalytic effects were strongly correlated for second shell mutants, with only a small number of outliers (Fig. 4*B*), and the correlation decreased substantially for the third shell residues and beyond (Fig. 4*C-D* and *SI Appendix*, Fig. S6 and Tables S4,S5). Unlike vanadate, however, many mutants in distal shells increased tungstate affinity, with the largest enhancements (10-fold) for mutations clustering near PafA’s distal Zn^2+^ ion (Fig. 4*E,F*).

**Fig. 4.**
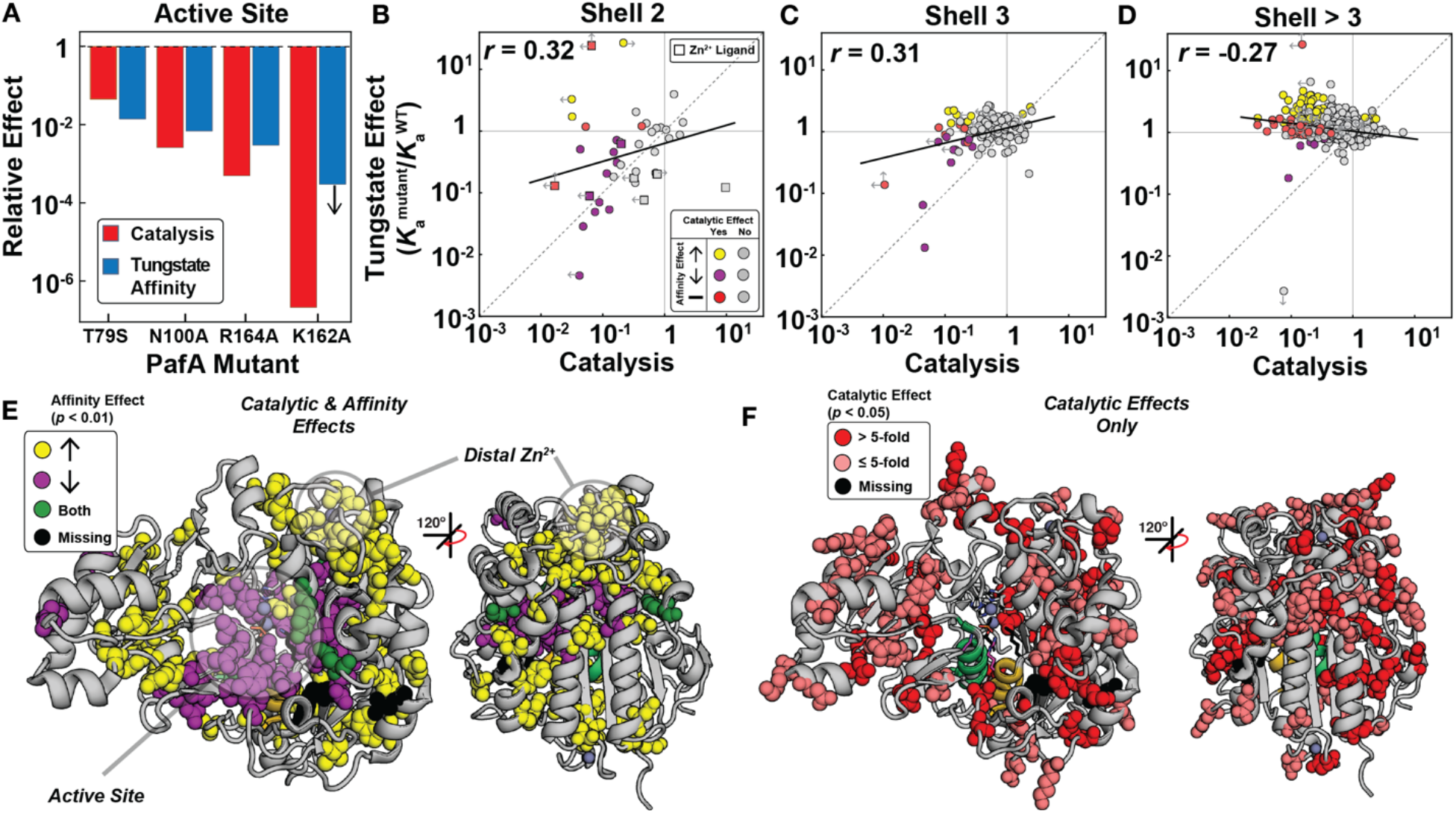
Effects of mutations throughout the PafA structure on tungstate binding and catalysis as a function of distance from the PafA active site. (**A**) Comparison of catalytic and tungstate affinity effects of PafA active site mutants, measured previously (24). (**B-D**) Comparison of catalytic and tungstate affinity effects for mutants of residues in the (**B**) second shell, (**C**) third shell, and (**D**) more distal shells. Labeled points in (**B**) denote mutants with significant catalytic but not affinity effects. (**E**) PafA structure showing locations of residues with significant catalytic and either deleterious (28 of 522 measurable residues) or enhanced (57 of 522 residues) tungstate affinity effects when mutated to either glycine or valine. (**F**) PafA structure showing positions in PafA that have significant catalytic but not significant affinity effects (80 of 522 residues). Catalytic data are from reference (23).

### Stronger correlation of catalysis with TSA affinity than ground state analog (GSD) affinity

A formal requirement for catalysis is the preferential stabilization of the transition state relative to the ground state. Consistent with this expectation, vanadate and tungstate binding to 1,004 PafA variants were more strongly correlated with catalysis than binding of Pi (a GSA and substrate for the reverse reaction) (Figs. 2*F,G* and *SI Appendix*, Fig. S7). This observed behavior is also consistent with the wealth of data demonstrating greater sensitivity of transition states (catalysis) than ground states (affinity) to mutations in many enzymes (e.g. (34–39)).

Vanadate and tungstate affinities also correlated well with one another (Fig. 5*A*, *r* = 0.53), especially for the active site and second shell mutants (*r* = 0.98 and *r* = 0.87, respectively), and better than each correlated against Pi affinity (Fig. 5*B,C*), as expected for a common covalently bound pentavalent (trigonal bipyramidal) species (25, 27).

**Fig. 5.**
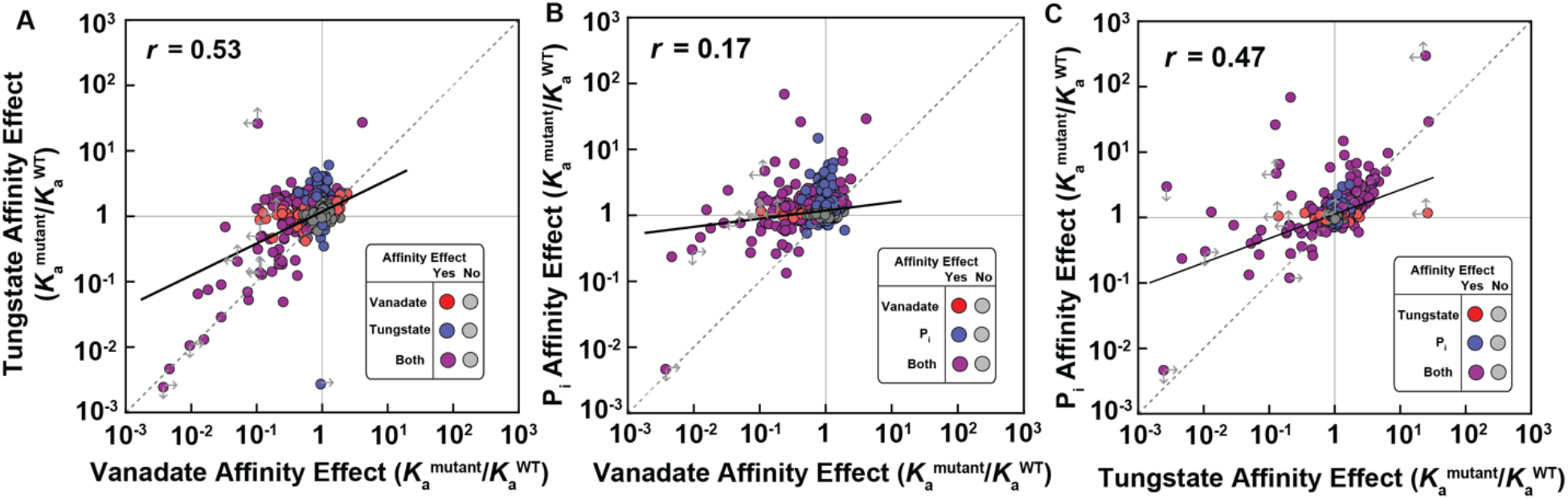
Comparison of measured binding affinities for transition state and ground state analogs. (**A**) Vanadate *vs*. tungstate affinity effects relative to WT (*K*_a_^mutant^/*K*_a_^WT^); (**B**) vanadate *vs*. Pi affinity effects; and (**C**) tungstate *vs*. Pi affinity effects. Gray markers have no significant effect on either of the ligands being compared; red, blue, and purple markers have significant effects as denoted in each legend. The Pi affinity data are from reference (23).

## Discussion

Our measurements of 1,944 affinity constants for vanadate and tungstate binding to 1,004 PafA variants combined with our prior data detailing impacts on catalysis and for Pi binding for these same mutants yield some expected results (23). The correspondence of catalytic (transition state, TS) and TSA effects in and around the active site matches the simplest expectation that direct interactions to TS or TSA atoms with similar charge and positioning contribute similar amounts of binding energy (Fig. 6*A,B*) and that removing groups that help position these contacting residues (second shell) yield similar (but smaller) energetic consequences (Fig. 6*C*). In the simplest case, this reduction in the magnitude of observed effects would report on how a mutation alters the fraction of time spent by an active site residue in a state that is competent for catalysis (Fig. 6*C*). For example, the active site arginine of *E. coli* alkaline phosphatase (R166) is mispositioned when either of its flanking aspartate residues are removed (D101 and D153), leading to losses of catalysis and TSA binding that are similar, but both smaller in magnitude than the effects from removal of the arginine itself (40, 41). Analogous positioning effects of non-active-site mutations on the positioning of active site residues have been observed in many other enzymes (e.g. (42–46)). However, other experimental outcomes were not anticipated.

**Fig. 6.**
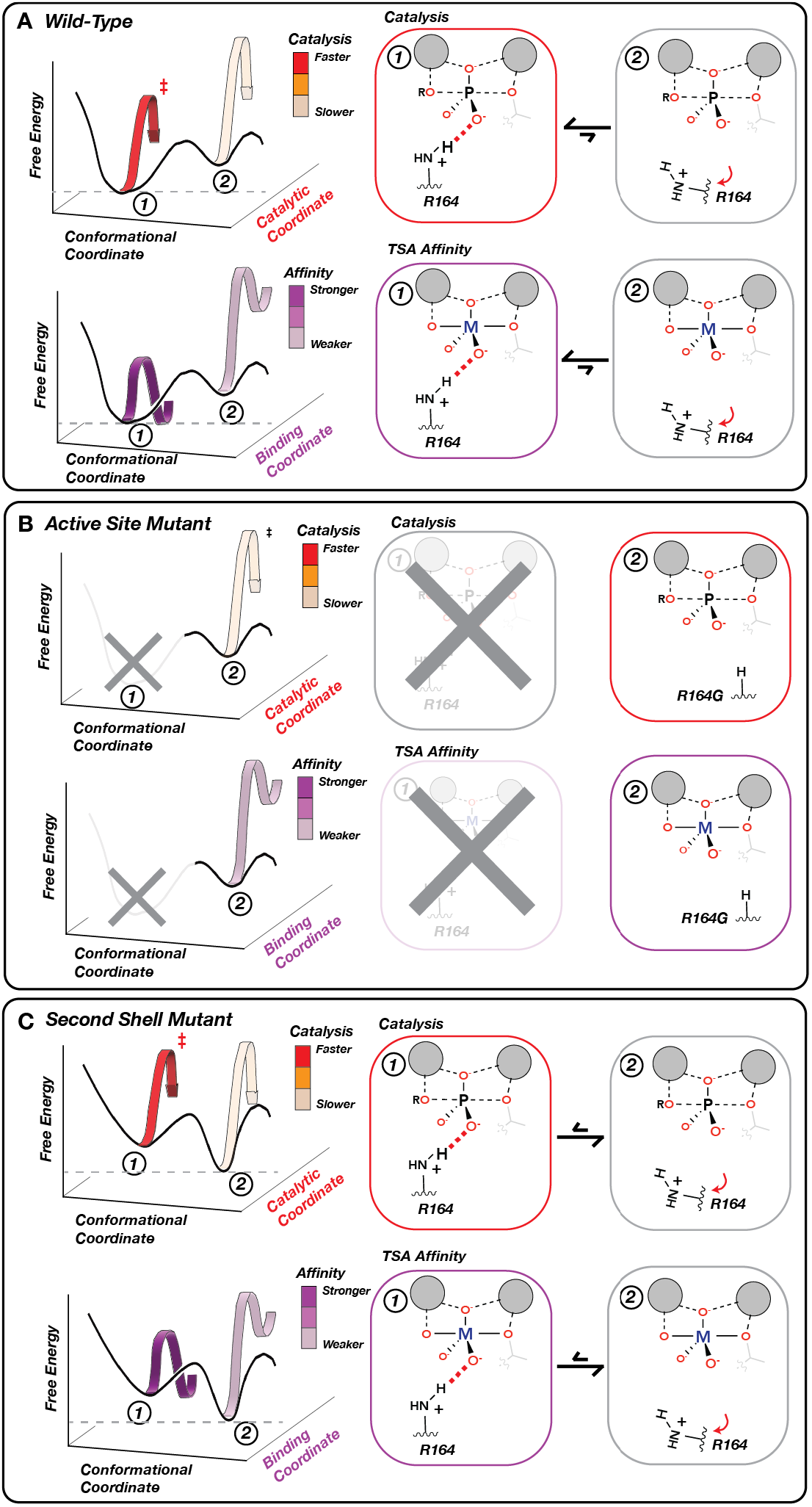
Physical models accounting for observed effects of mutating active site and second shell residues. **(A)** A WT PafA active site residue shown in a conformational equilibrium between two states. In State 1 (favored), R164 donates a hydrogen bond to an oxygen atom in the TS and TSA, promoting catalysis and TSA binding (M = V, W). In State 2 (disfavored), R164 is mispositioned so that the hydrogen bond no longer forms, reducing both catalysis and TSA binding. **(B)** Ablation of the R164 side chain eliminates State 1, reducing catalysis and TSA binding to the same extent (to the level of WT State 2 alone (**A**)). **(C)** Mutating a second shell residue that positions R164 destabilizes State 1 and increases occupancy of State 2. This change in the conformational equilibrium equally reduces catalysis and TSA binding, proportional to the increased occupancy of State 2.

### Why does the TSA–TS correspondence dissipate for more distal mutations?

TS and TSA effects diverge with beyond the second shell, with mutations that are deleterious to catalysis no longer impairing vanadate or tungstate binding (Figs. 3*A-D*, 4*A-D* and *SI Appendix*, Figs. S5, S6). While the observation of widespread effects of distal mutations on chemical catalysis indicates that those positions communicate to the active site, this divergence is not accounted for by the simple model depicted in Fig. 6. Instead, we propose that alterations to peripheral residues can be transmitted to the active site through a series of more subtle conformational adjustments, analogous to previously described mechanisms communicating allosteric effects (47–49). As all proteins exist as an ensemble of (many) conformations whose occupancies are determined by the protein’s free energy landscape (50–52), mutations that impact catalysis but not TSA binding could increase relative occupancies of microstates that are less catalytically potent but not compromised in TSA binding (Fig. 7*A,B*). Physically, this divergence likely stems from differences in size, geometry and/or charge distribution between TSAs and the actual transition state. Vanadate and tungstate are larger than phosphate, with mean bond lengths of 1.8 Å for vanadate and 1.7 Å for tungstate (compared to 1.5 Å for phosphate) (*SI Appendix*, Fig. S8).

**Fig. 7.**
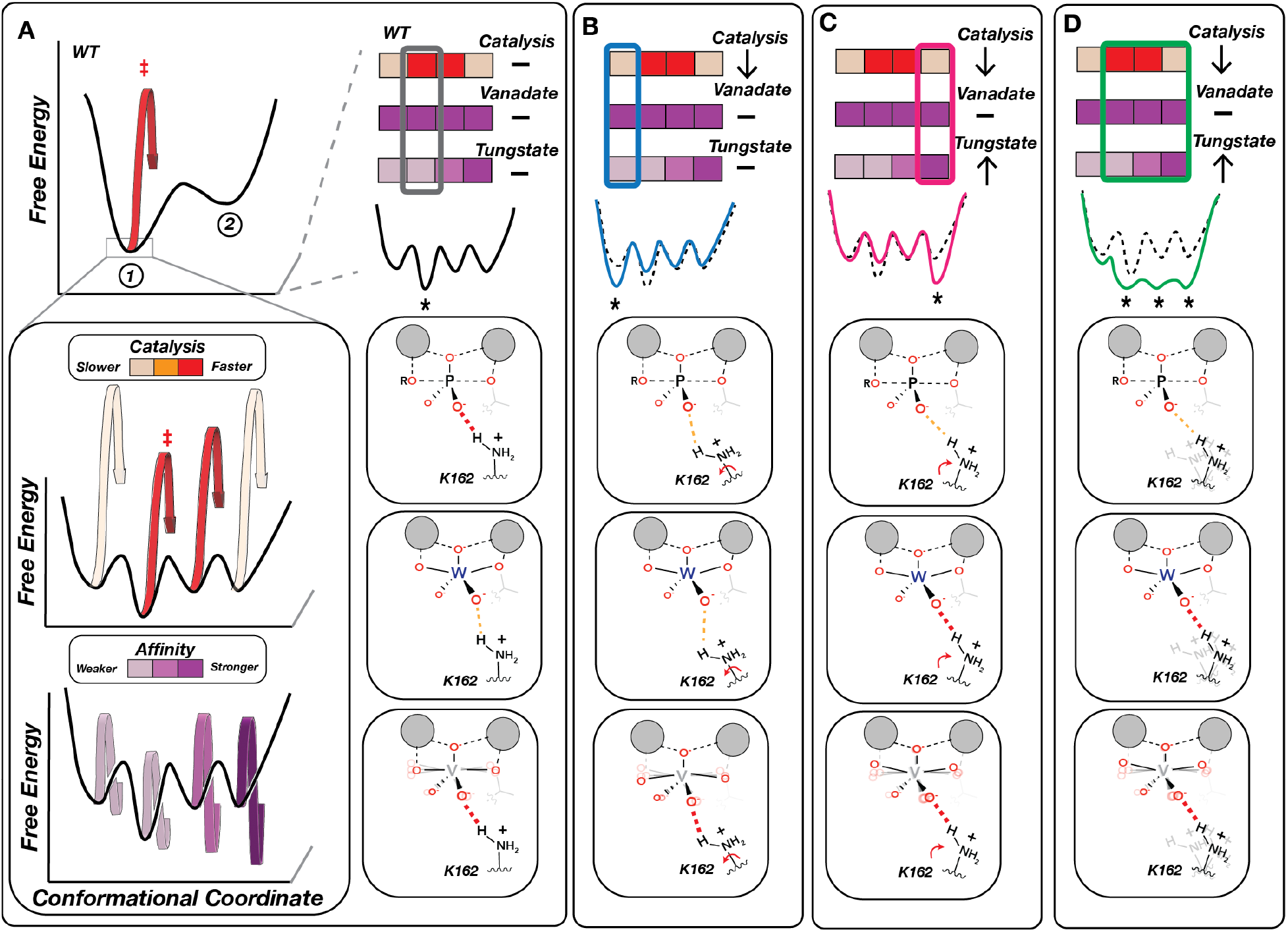
Physical models for the impacts of distal residue mutations that decouple catalytic and TSA affinity effects. **(A)** WT-favored State 1 is itself an ensemble of microstates (that is, states with smaller conformational changes than illustrated for the mutants in Fig. 6) with different levels of catalysis (middle panel) and TSA binding (bottom panel), denoted by color bars. The most populated WT microstate is optimal for catalysis but not for tungstate binding, whereas vanadate’s conformational flexibility allows the ligand to adopt more optimal geometries in each microstate. (B-D) Examples of distal mutational effects with differential impacts on catalysis and TSA binding due to alternation in microstate distributions. **(B)** Stabilization of a less catalytically-competent microstate that maintains WT TSA affinities (blue). **(C)** Stabilization of a less catalytically-competent microstate that preferentially binds the distorted tungstate geometry (pink). **(D)** Flattening of the conformational landscape due to increased flexibility results in similar occupancies of multiple microstates, including some with increased affinity for the distorted geometry of tungstate (green). Not shown are distal effects that phenocopy the effects described in Figure 6B.

### Why do many distal mutants <u>increase</u> tungstate affinity?

Most mutations that increase tungstate affinity are far from the active site (149 of 155 are in the third shell or beyond; *SI Appendix*, Tables S4 and S5), and at least 62 of the 149 are also *deleterious* for catalysis (*SI Appendix*, Fig. S9). These results, while not anticipated, are readily accounted for from an ensemble perspective in which distal substitutions shift the distribution to increase occupancy of microstates that are less catalytically proficient and bind tungstate more strongly (Fig. 7*C*). The observation that many more glycine than valine mutations increase tungstate binding (95 *vs*. 50, *SI Appendix*, Tables S6 and S7 and *SI Appendix*, Figs. S10 and S11) supports this model and the notion that distal residues act to restrict PafA’s conformational ensemble (Fig. 7*D*). The fact that distal mutants can *increase* binding of a TSA that differs from the actual transition state by tenths of an Ångström suggests that distal residues play a critical role in precisely positioning active site residues.

Structural data further support this model and the high degree of positioning. Tungstate bound to the related *E. coli* alkaline phosphatase has a 164° axial O-W-O bond angle, more acute than expected for a pentavalent trigonal bipyramidal species (*SI Appendix*, Fig. S12) but consistent with small-molecule tungsten compounds (*SI Appendix*, Fig. S13*A,B*). This difference suggests a suboptimal fit for the larger and more bent tungstate in sites evolved for the phosphoryl transition state and that distal mutations may alter the active site to better accommodate tungstate (Fig. 7*C,D*).

### Why do mutations that increase tungstate affinity not increase vanadate affinity?

We observed increased binding of tungstate but not vanadate from distal mutations (Fig. 2*F,G*). Two models, based on the different properties of pentavalent vanadate and tungstate, could account for this difference. The axial bond angle is more acute for tungstate than vanadate (*SI Appendix*, Fig. S12) (25, 27); thus, tungstate may be more destabilized so that subtle conformational alterations partially relieve this destabilization. Alternatively, there appears to be considerable flexibility in pentavalent O–V–O_axial_ bond angles compared to phosphate and tungstate (*SI Appendix*, Fig. S13*E,F*); this flexibility may reflect a broader and shallower potential well for vanadate so that there is no significant binding advantage from a broadened enzyme ensemble (Fig. 7*D*).

#### Implications and Future Directions

There is now extensive evidence for widespread allostery at many surface sites of proteins and enzymes (48, 51, 53), and directed evolution experiments often uncover distal mutations that increase the rate of the target reaction (e.g., (54–59)). Our prior results revealed communication from many distal residues to PafA’s active site that impact catalysis (23). Here, our investigations of TSA binding revealed that distal residues throughout PafA shape the conformational landscape of the PafA active site on the tenth-Ångström length scale. As expected, residues in and directly contacting the active site are deleterious to catalysis and TSA binding. But as one moves away from the active site, deleterious impacts on catalysis remain common while those on TSA binding are diminished or eliminated, and tungstate binding can even be enhanced (Movies S1-S3).

The intricate and broadly distributed enzyme features involved in shaping an enzyme’s conformational landscape to optimize catalysis may complicate the design of highly efficient enzymes. Our results suggest that the enzyme beyond the active site and second shell is integral to this optimization (Movies S1-S3). However, these effects are subtle and thus difficult to accurately model. The need to consider structure as an ensemble of states rather than a singular structure further confounds the development of accurate predictive models for enzyme design.

Our results suggest that distal residues communicate to the active site by necessity to optimize function. If this is the case, surface mutations near them could “create” binding sites that affect the distal residue’s conformational preferences and thereby alter catalysis. According to this view, the involvement of intermediate shells in optimizing active site conformational preferences enables surface residues to influence their properties. Moreover, the widespread incidence of distal effects suggests that allostery is a highly evolvable trait, providing a biophysical rationale for the prevalence of allostery (60, 61).

High-throughput enzymology results underscore the complexity of “function” and the value of and need to measure multiple kinetic and thermodynamic parameters (*e.g*. (23) & this work). Mutational effects on catalysis are widespread, extending to the enzyme’s surface with unanticipated patterns (Movies S1-S3), and different substitutions to the same residue can have different effects and impact different reaction steps (23). Applying standard dimensionality reduction approaches to the multi-parameter PafA mutational data reveals a complex pattern without distinct and well-separated clusters, further underscoring this complexity (*SI Appendix*, Fig. S14). More sophisticated clustering and dimensionality reduction techniques that consider three-dimensional spatial relationships may help reveal common physical features that alter particular aspects of catalysis. Nevertheless, to understand how natural enzymes achieve their enormous rate enhancements and exquisite specificities and to reproduce these properties in designed enzymes will likely require physical and energetic models that extend beyond dimensionality reduction approaches. Providing testable quantitative predictions and models will likely involve integrating free energy landscapes and conformational ensembles with sufficiently accurate models of the local forces at play in and around enzyme active sites. Given the enormity of this challenge, it is likely that HT-MEK and other quantitative high-throughput approaches will be indispensable in generating ground truth data for model generation and for robustly testing predictions made by current and future models.

## Materials and Methods

### On-chip expression, HT-MEK device set up, and expression and purification of PafA mutants

HT-MEK devices were used to recombinantly express PafA mutants as described previously (23). Briefly, devices were fabricated using standard soft lithography techniques and then aligned to PafA-eGFP mutant plasmid arrays deposited on epoxy-coated slides using a custom microarray printer. Following alignment, device surfaces were specifically patterned with anti-GFP antibody beneath ‘button’ valves and passivated with BSA elsewhere; after surface patterning, cell-free expression mix was introduced into each chamber, solubilizing printed plasmid DNA. PafA-eGFP mutants were expressed at 37°C for 45 min, incubated at room temperature for 90 min, and then immobilized on the surface of anti-GFP patterned ‘button’ valves and purified via washing with ‘button’ valves closed.

### Measuring vanadate and tungstate affinities using HT-MEK

Inhibition constants (*K*_i_) for vanadate and tungstate were determined via competitive inhibition assays with the fluorogenic substrate cMUP (50 μM) as previously described for measurements of inorganic phosphate (Pi) inhibition (23). Vanadate measurements used concentrations of 0, 0.25, 0.5, 1, 2, 5, and 10 μM for vanadate, in some cases with 100 μM vanadate added to the series; tungstate measurements used concentrations of 0, 0.5, 1, 2, 5, 10, 25, 100, 250, 500 and 1000 μM for all experiments except two, which went up to 100 μM and 500 μM, respectively. For each mutant replicate, we fit initial rates of cMUP hydrolysis at each inhibitor concentration and then fit these initial rates to a standard competitive inhibition model, as previously described (23).

To ensure accurate quantification of TSA affinities (*K*_D_), all experiments used a substrate concentration (50 μM) that is below the previously-measured cMUP *K*_M_ (23) for nearly all (>90%) of PafA variants and provides high signal to noise in fluorescence time-course measurements. As observed *K*_i_ values are proportional to both the substrate *K*_M_ and the intrinsic affinity (*K*_D_), we used the Cheng-Prussoff relationship (62) and previously measured *K*_M_ values for cMUP hydrolysis for each mutant (23) to correct for these modest systematic differences between the measured *K*_i_ and the *K*_D_. Returned *K*_D_ values were within two-fold of the measured *K*_i_ values for 88% of mutants (*SI Appendix*, Fig. S15).

We used multiple quality control metrics to ensure that measurements from each chamber were due to the mutant printed in that chamber and not from cross-contamination between chambers, as described previously (23). To ensure each mutant expressed and immobilized successfully, we manually confirmed that each chamber contained a measurable eGFP spot (corresponding to [E] > 0.3 nM) and that each spot was free of fluorescence artifacts. To ensure high quality activity measurements, we included a catalytically inactive ‘fiducial’ mutant T79G or chamber lacking a mutant DNA template (“Skipped”) in every 7^th^ chamber, considered measured rates within these chambers to represent a local ‘background’ rate, and culled data from any chambers in which measured activity was less than 5-fold greater than an interpolated local background rate. To further ensure accurate activity measurements for the most catalytically-compromised mutants, we implemented a tiered measurement strategy in which all mutants within the glycine and valine scanning libraries were measured together initially and then the slowest mutants were selected, reprinted, and assayed again separately.

As described previously for Pi (23), we first normalized fitted *K*_D_ values for each mutant to the median *K*_D_ of the wild-type replicates within a given experiment and then calculated the median values for each mutant across experiments. *K*_D_ values from experiments containing only the slowest mutants (which did not include wild-type PafA) were not normalized. We determined the statistical significance of *K*_D_ effects for each mutant relative to WT PafA using bootstrap hypothesis testing (*p* < 0.01) as previously described (23).

All progress curves, initial rate fits, inhibition curves, fitted *K*_i_ values, and *K*_D_ values after correction for each individual replicate measurement of each mutant, as well as median *K*_D_ values and measures of statistical significance for each mutant, are available in CSV and PDF files in the associated Open Science Foundation (OSF) repository (https://osf.io/k8uer/).

### Identifying upper and lower *K*_D_ limits

Upper and lower *K*_D_ limits arise when the measured (apparent) *K*_i_ is less than the lowest concentration of inhibitor assayed or greater than the highest concentration of inhibitor assayed, respectively. We flagged *K*_D_ values as lower limits if these values were greater than two-fold above the highest vanadate or tungstate concentration for >50% of all replicates of that mutant, as previously implemented for Pi affinity measurements (23). *K*_D_ values were flagged as upper limits if the measured *K*_i_ was two-fold lower than the lowest non-zero inhibitor concentration. Upper limits can also arise if the measured cMUP *K*_M_ values are upper limits, as this can introduce error in the Cheng-Prussoff correction; for these mutants the *K*_D_ values were denoted as limits. Upper and lower limits are denoted by arrows in the main text and supplementary figures.

### Assignment of residues to active site interaction shells

To determine the minimum number of interactions between each residue and the active site and assign residues to interaction shells, we used GetContacts (https://getcontacts.github.io) to identify all contacts between residues in the WT PafA crystal structure. We then defined the active site as composed of all residues or ions making direct contacts with the substrate (T79, N100, K162, and R164, and the two Zn^2+^s.) The set of residues making at least one contact to any of these active site components was then defined as the second shell. Residues making at least one contact to any of the second shell residues but not contacting the active site were defined as third shell residues. Residues were then successively assigned to the more distal shells in the following general manner: for shell *N*, residues were defined as residues making contacts to shell *N*-1, but not to any of the residues in the lower shells.

### Multi-parameter data visualization using UMAP

UMAP (63) was performed using its implementation in skikit-learn (64) in a Jupyter notebook (available at the OSF repository). The catalytic (*k*_cat_/*K*_M,chem_.) effects and vanadate, tungstate, and Pi affinities (log_10_ transformed) for all glycine and valine mutants for which these parameters were experimentally measurable (*n* = 928) were imported and the UMAP reducer was trained and the two-dimensional dimensionality-reduced array was output using the “reducer.fit_transform” function.

### Obtaining geometric parameters of vanadium, tungsten, and phosphorus compounds from the Cambridge Structural Database (CSD)

We used the CSD Python API to search for all molecules containing at least one M-O bond, where M denotes vanadium, tungsten, or phosphorus. We then used the GeometryAnalyzer function to determine the number of bonds to the M atom, identify each M-O bond, assess if the oxygen was esterified or non-esterified, and determine all M-O bond lengths and bond angles. We then classified the valency of each molecule based on the number of bonds to the M atoms, and classified its geometry (tetrahedral, square pyramidal/planer/octahedral, trigonal bipyramidal, or undefined) based on bond angle values, as defined previously (25).

For the comparisons of tetrahedral forms, we selected all molecules containing only M-O oxides. We also included M-O oxide distances measured previously in the literature but not deposited in the CSD (65, 66). For the comparisons of tetrahedral esters, we selected all molecules containing a mixture of oxides and M-O-C esters and compared the bond length distributions of each type. As the number of esterifying atoms on each oxygen strongly affects M-O bond lengths, we only considered singly esterified oxygens when comparing lengths of esterified M-O bonds.

For the comparisons of pentavalent molecules, the total number of molecules was limited, (especially for tungsten compounds). We therefore selected all pentavalent molecules containing a mixture of oxide bonds and at least one M-O-C ester, again only comparing oxides or singly esterified oxygens.

All code used to download, parse, and analyze these data is available at the OSF repository (https://osf.io/k8uer/).

### Geometric parameters of vanadium and tungsten compounds bound to proteins in the PDB

PDB structures containing tungstate and vanadate compounds bound to proteins were analyzed using the Bio.PDB package in Python. To infer connectivity and geometry of the bound molecules in each structure, we first identified all neighboring atoms within 2.5 Å of the metal atom. For each neighboring oxygen atom, we then calculated the number of atoms bonded to that atom within 2.5 Å. We assigned oxygens forming only one bond as oxides and others as esters and then calculated all bond lengths and angles involving the metal atom. Structures with five atoms bonded to the metal atom and one (and only one) bond angle larger than 160 degrees (the angle between axial atoms) were defined as having pentavalent trigonal bipyramidal geometry; structures with four atoms bonded to the metal atoms (based on the above criteria) were assigned as tetrahedral.

## Data, Materials, and Software Availability

All experimental data are provided as a CSV file in Supplementary Data. Additionally, all experimental data, including per-experiment and per-mutant summary reports of the on-chip measurements, as well as all code used to process the data are available in the Open Science Foundation repository (https://osf.io/k8uer/).

## Acknowledgements

We thank Hiwot Anteneh for technical assistance and helpful discussions, as well as all members of the Herschlag and Fordyce laboratories for helpful discussions. This work was supported by NIH grant R01 (GM064798) awarded to D.H. and P.M.F., an Ono Pharma Foundation Breakthrough Innovation Prize, and the Gordon and Betty Moore Foundation (grant 8415). P.M.F. acknowledges the support of an Alfred P. Sloan Foundation fellowship and is a Chan Zuckerberg Biohub Investigator. C.J.M. acknowledges the support of a Canadian Institutes of Health Research (CIHR) Postdoctoral Fellowship. D.A.M. acknowledges support for the Stanford Medical Scientist Training Program and a Stanford Interdisciplinary Graduate Fellowship (Anonymous Donor).

