## Supplementary material for "High-throughput enzymology reveals mutations throughout a phosphatase that decouple catalysis and transition state analog affinity": SI Appendix

PNAS

**This PDF file includes:**

Figures S1 to S15
Tables S1 to S9
Legends for Movies S1 to S3

**Other supporting materials for this manuscript include the following:**

Movies S1 to S3
Dataset S1

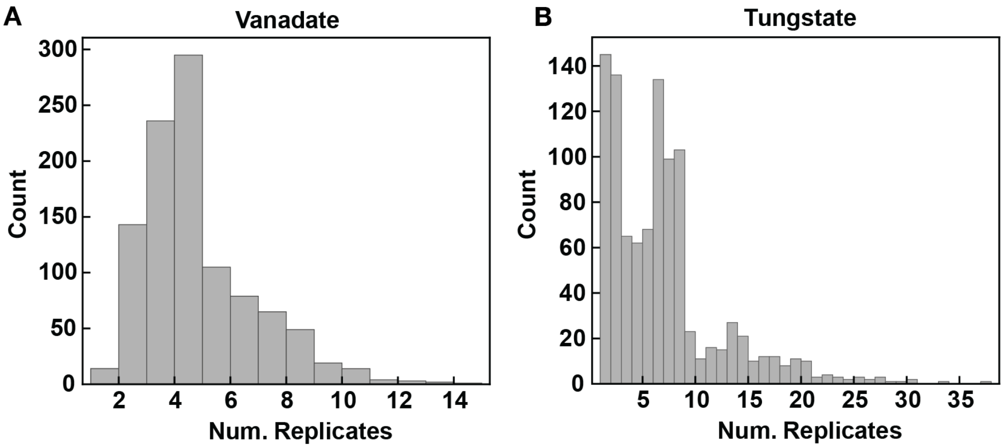


**Fig. S1.** Number of replicate affinity measurements for (A) vanadate and (B) tungstate for each mutant from the glycine and valine scanning libraries.

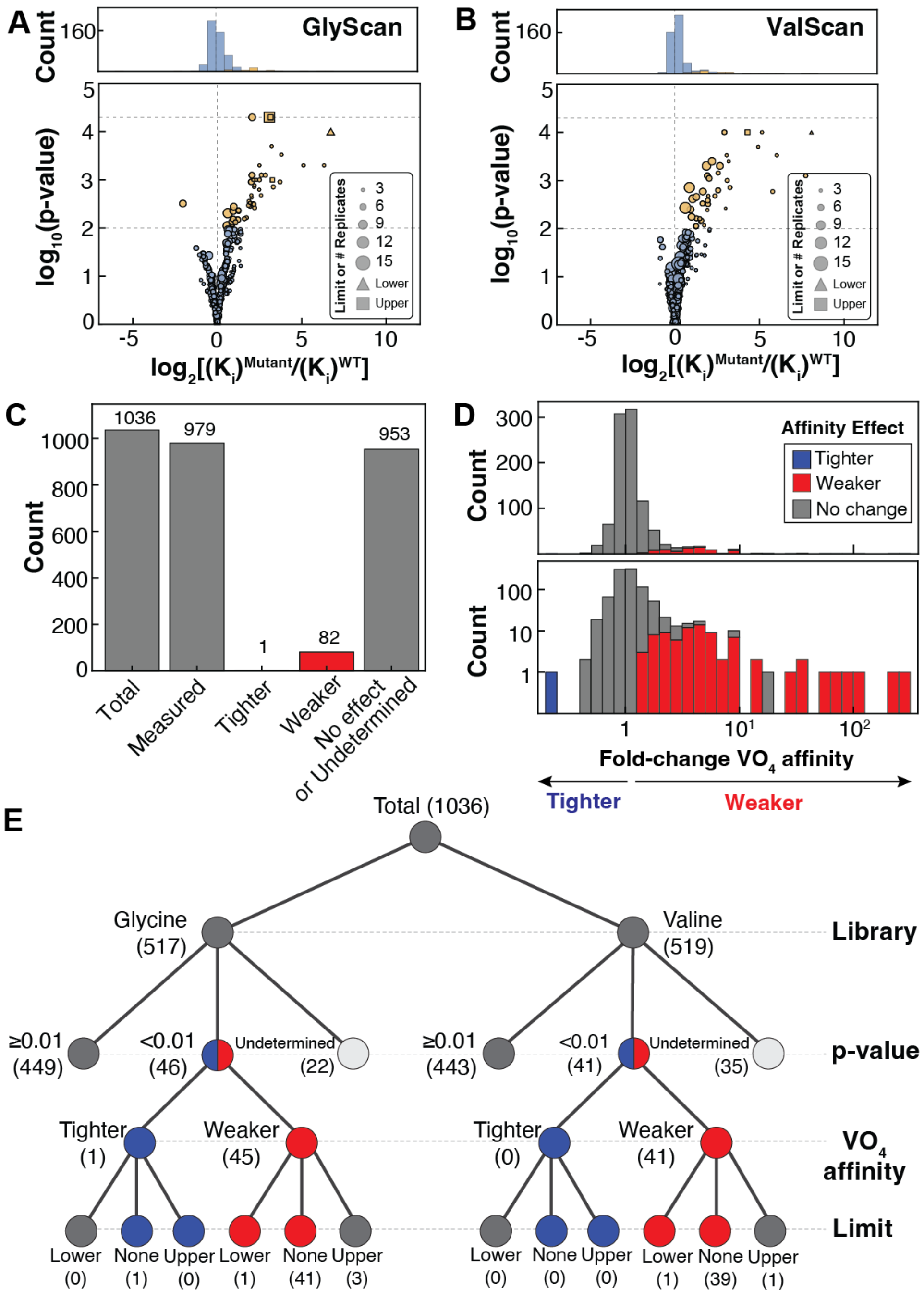


**Fig. S2**. Measured effects of mutations on vanadate ($VO_4$) affinity for all mutants within the glycine and valine scanning libraries. (A) Volcano plot of vanadate affinity effects for all glycine mutants. (B) Volcano plot of vanadate affinity effects for all valine mutants. (C) Number of mutants for which vanadate affinity was measured, and number of mutants with changes in vanadate affinity ($p<0.01$), and with unchanged (WT-like) vanadate affinity ($p\geq0.01$). (D) Distribution of measured vanadate affinity effects in both libraries, in linear (top) and logarithmic scale (bottom). (E) Tree diagram summarizing the number of mutants with tighter, weaker, and unchanged vanadate affinity in the glycine and valine scanning libraries.

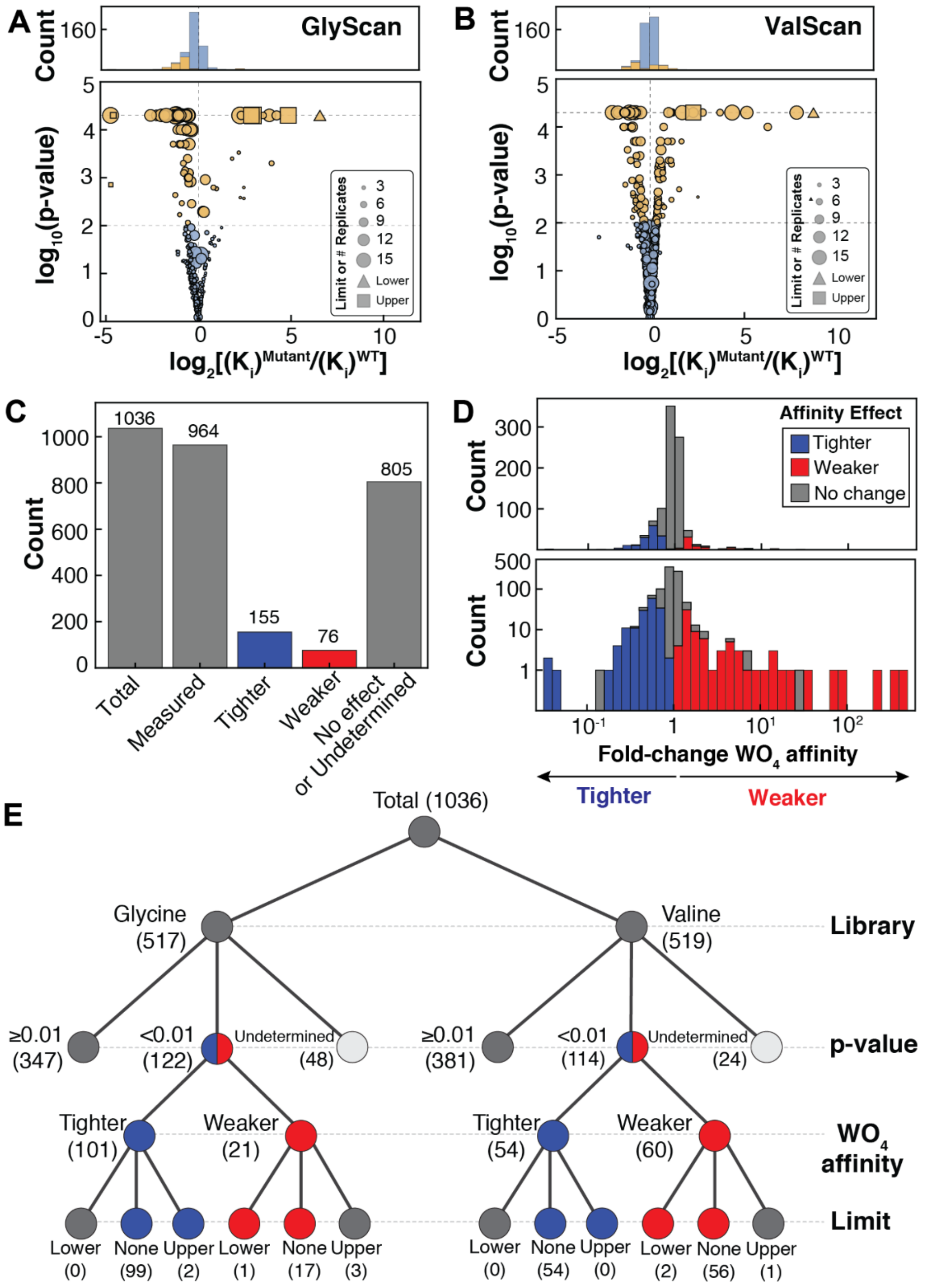


**Fig. S3**. Mutational effects on tungstate ($WO_4$) affinity within the glycine and valine scanning libraries. (A) Volcano plot of tungstate affinity effects for the glycine scanning library. (B) Volcano plot of tungstate affinity effects for the valine scanning library. (C) Number of mutants for which tungstate affinity was measured, and number of mutants with changes in tungstate affinity ($p<0.01$), and with unchanged (WT-like) tungstate affinity ($p\geq0.01$). (D) Distribution of measured tungstate affinity effects in both libraries, in linear (top) and logarithmic scale (bottom). (E) Tree diagram summarizing the number of mutants with tighter, weaker, and unchanged tungstate affinity in the glycine and valine scanning libraries.

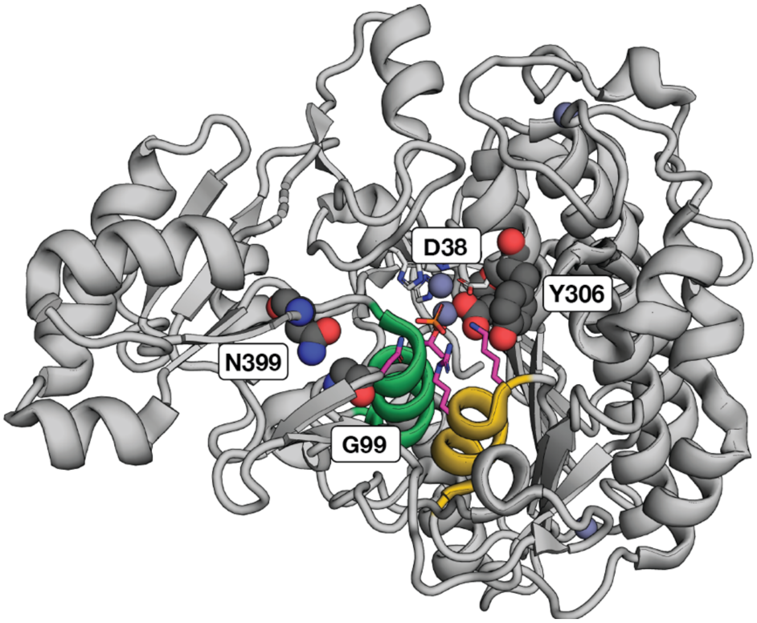


**Fig. S4.** Location of the four second shell mutants (spheres) preferentially influencing catalysis compared to vanadate affinity on the PafA structure.

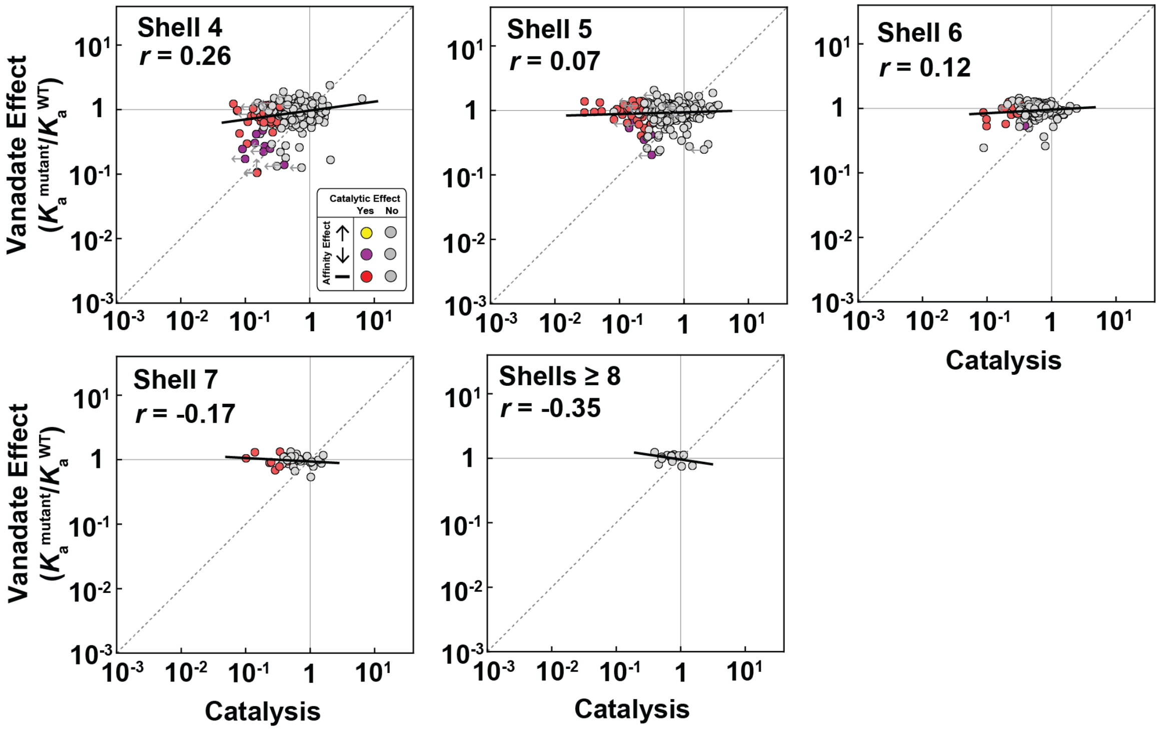


**Fig. S5**. Comparison of vanadate affinity and catalytic effects by shell, for each individual shell beyond the second shell (also see **Fig. 3A-D**). Gray dashed lines denote 1:1 line, and solid black lines are the best fit lines.

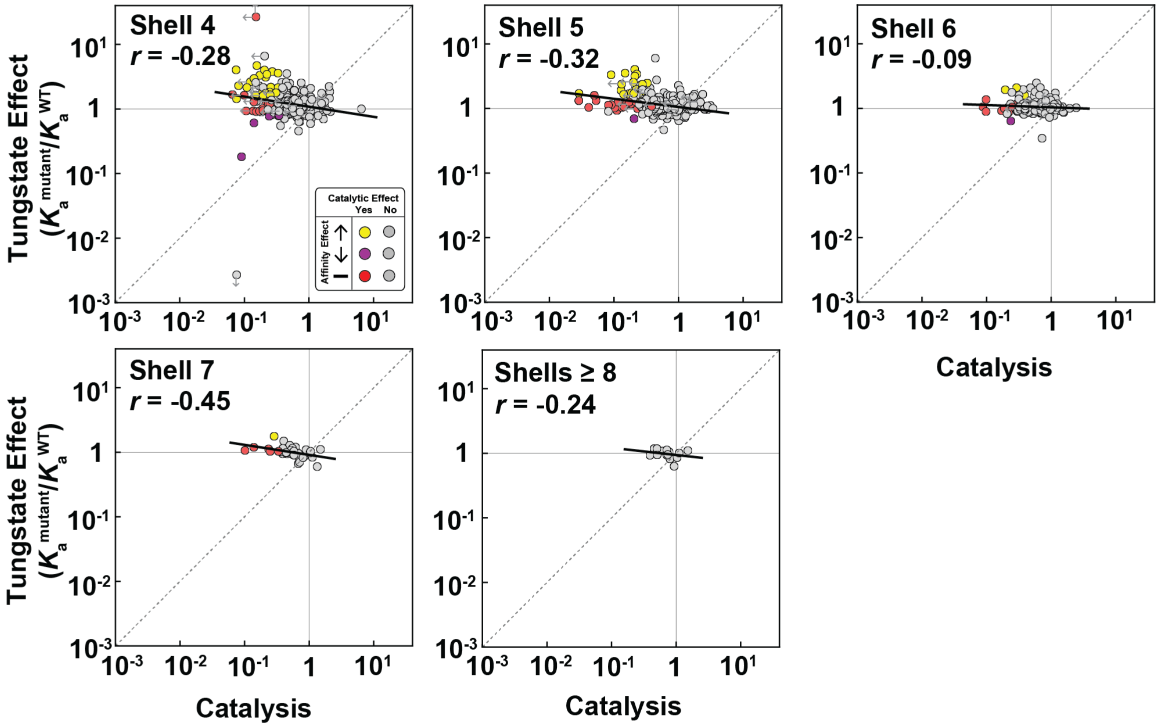


**Fig. S6**. Comparison of tungstate affinity and catalytic effects by shell, for each individual shell beyond the second shell (also see **Fig. 4A-D**). Gray dashed lines denote 1:1 line, and solid black lines are the best fit lines.

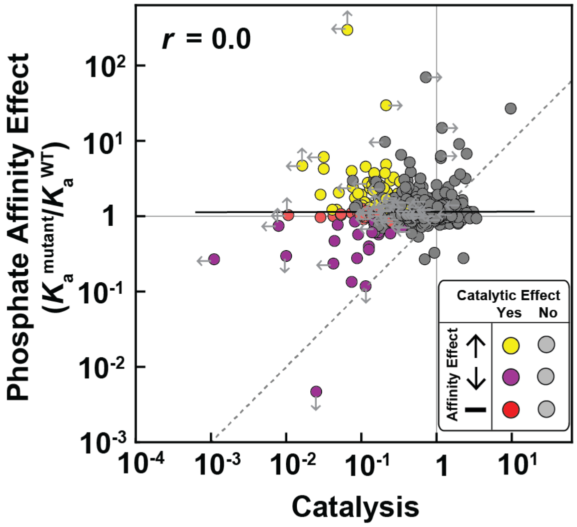


**Fig. S7.** Comparison of catalytic effects and previously-measured $P_i$ affinity effects (1), colored by significance of effects.

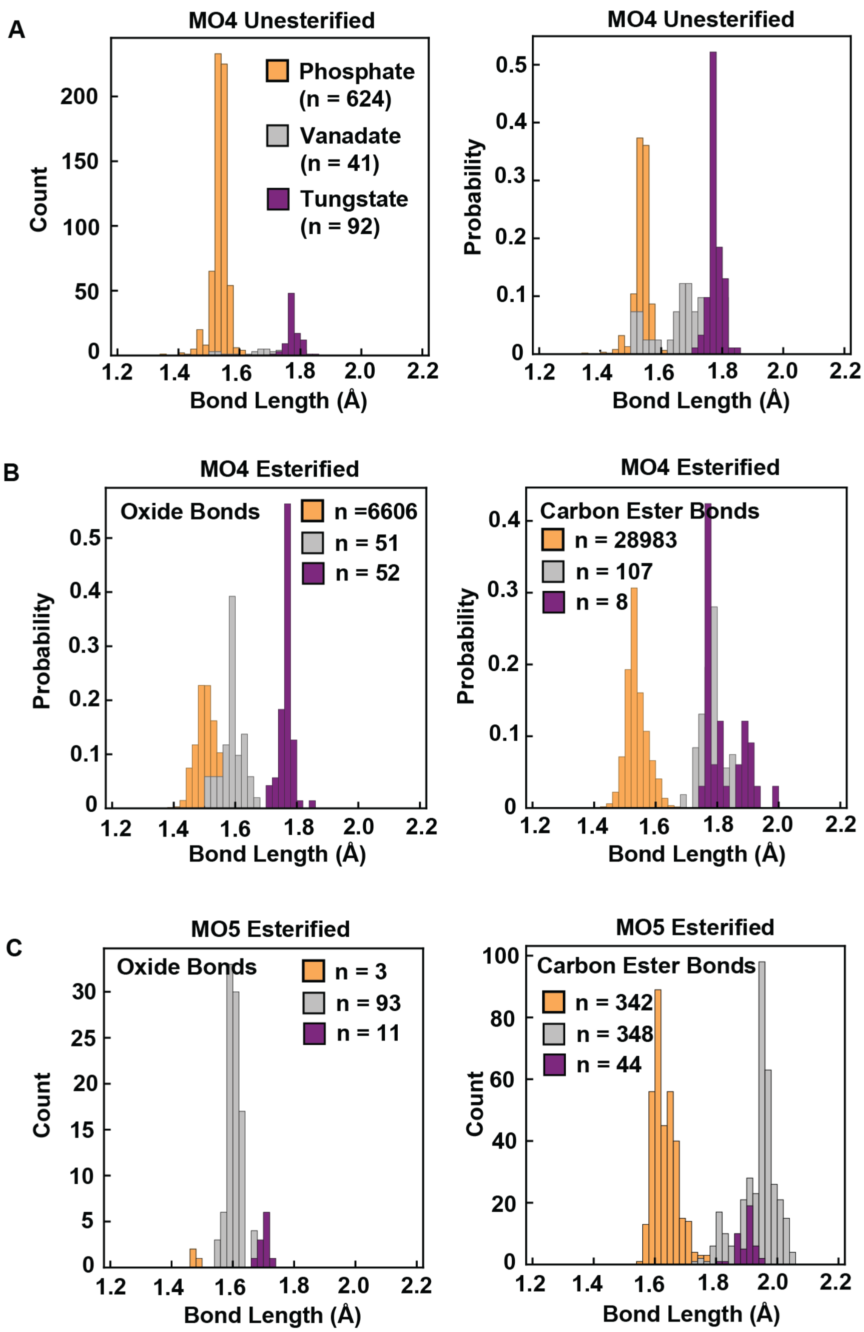


**Fig. S8.** (A) Unesterified M-O bond lengths for fully unesterified tetrahedral $MO_4$ phosphate, vanadate, and tungstate compounds in the CSD and literature (2, 3). Numbers in the legend denote number of unique bond lengths for each set. (B) Unesterified M-O oxide (left) and esterified M-O-C (right) bond lengths for $MO_4$ phosphate, vanadate, and tungstate compounds containing carbon esters in the CSD. Distributions are shown as probability densities as the number of phosphorus compounds far exceeds the numbers of vanadium and tungsten structures. (C) Unesterified M-O oxide (left) and esterified M-O-C (right) bond lengths for pentavalent ($MO_5$) phosphorus, vanadium, and tungsten compounds containing carbon esters in the CSD.

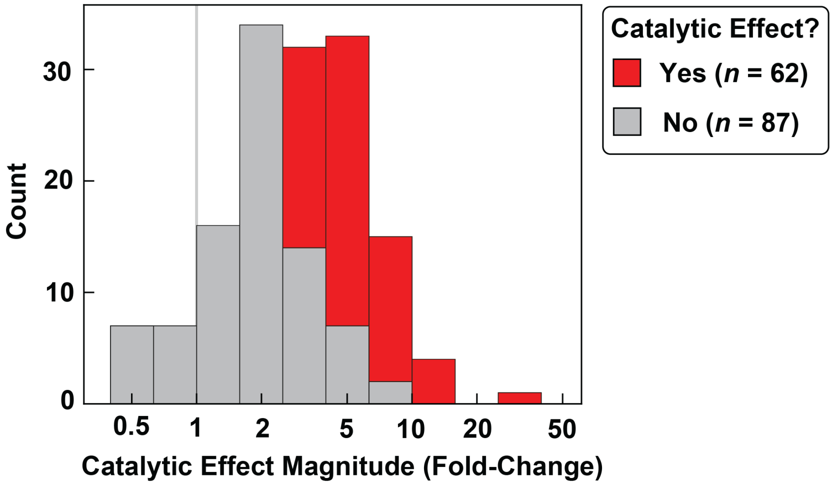


**Fig. S9.** Catalytic effect magnitudes (deleterious fold-change) for mutants with increased tungstate binding beyond the second shell (stacked histogram).

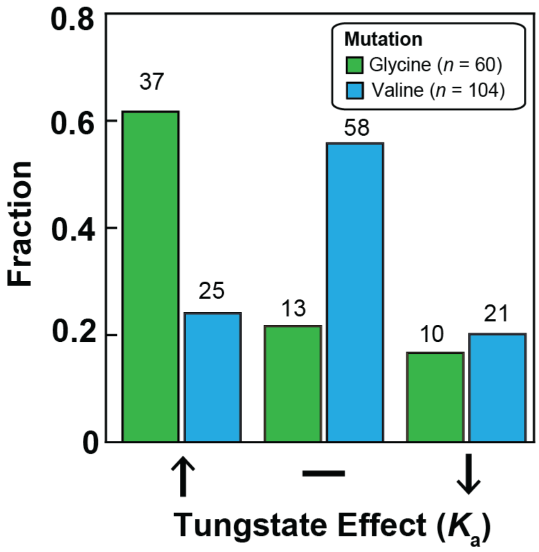


**Fig. S10.** Fractions of the total set of glycine and valine mutants with significant deleterious catalytic effects that have increased, unchanged, or decreased tungstate affinity. Bars are labeled by the total number of mutants within each set.

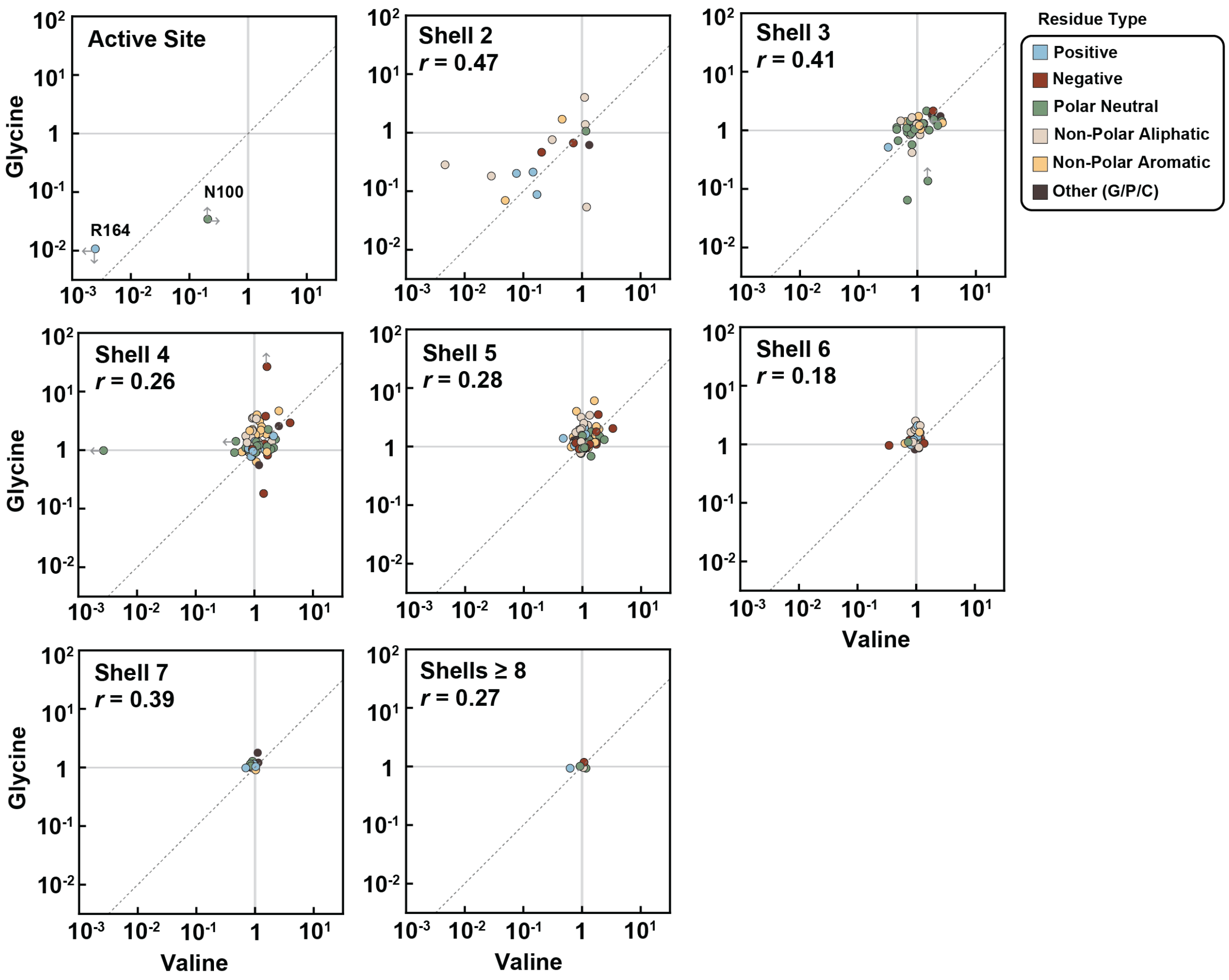


**Fig. S11.** Comparison of glycine and valine substitution effects on tungstate affinity within each shell. Points are colored by the chemical property of the wild-type residue, as indicated by the legend. For the active site mutant comparison, only mutants of N100 and R164A were measurable, as rates of hydrolysis of the glycine and valine substitutions of T79 and K162 were below the limit of detection.

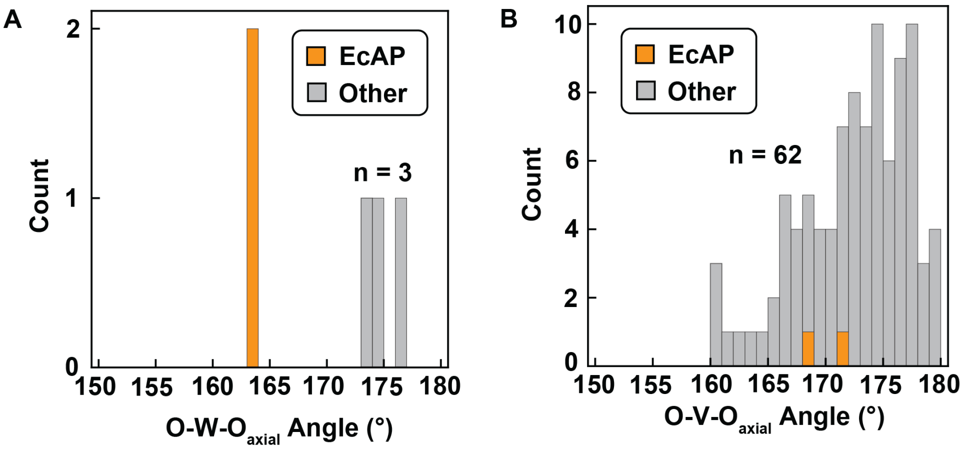


**Fig. S12.** Distributions of axial O-M-O angles for (A) tungstate and (B) vanadate bound to enzymes in the PDB and comparisons with axial angles observed for *E. coli* alkaline phosphatase (EcAP) bound to tungstate and vanadate.

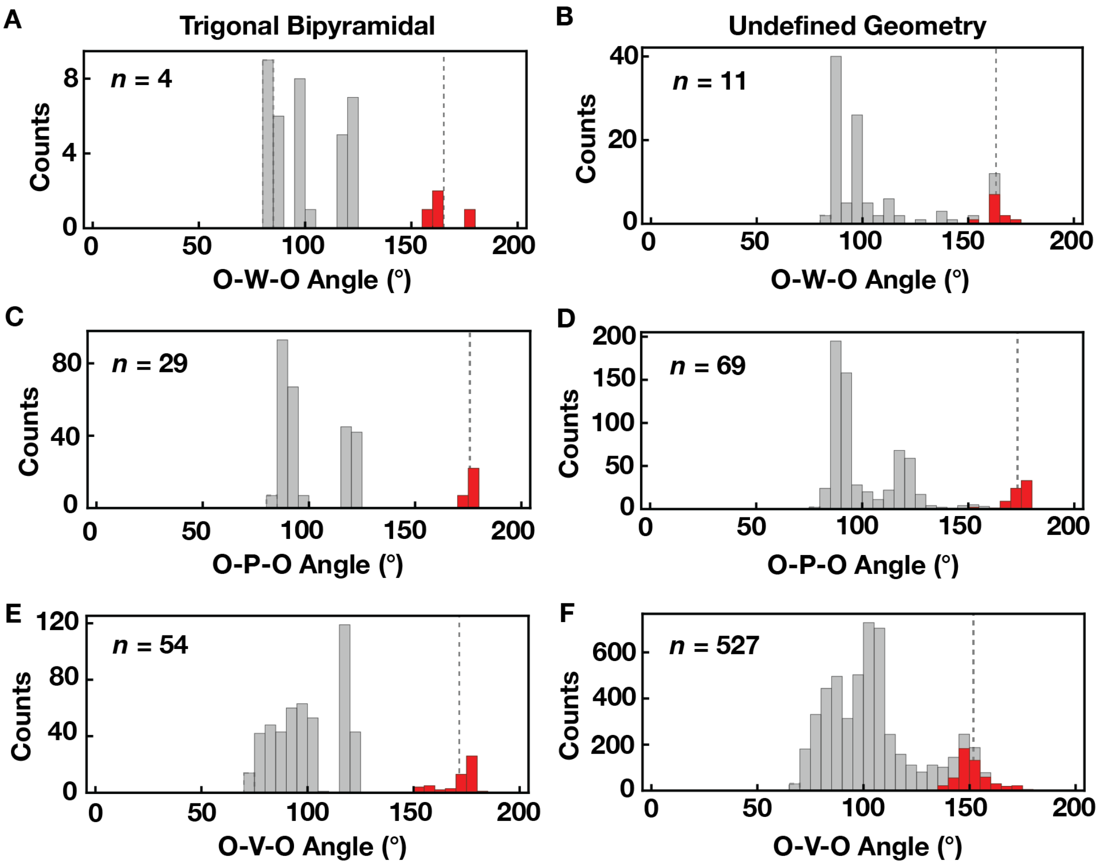


**Fig. S13.** Bond angle distributions of small molecule pentavalent tungsten, phosphorus, and vanadium structures. (A, B) Distribution of bond angles for trigonal bipyramidal tungsten (A) compounds and compounds for which the geometry cannot be unambiguously assigned ("undefined") (B) (see Methods). (C, D) Distribution of bond angles for trigonal bipyramidal phosphorus compounds and phosphorus compounds with undefined geometry. (E, F) Distribution of bond angles for trigonal bipyramidal vanadium compounds and vanadium compounds with undefined geometry. For the trigonal bipyramidal geometries, red and gray bars denote the axial O-M-O angles and all other angles, respectively, and for the undefined geometries red denotes the largest angle for each compound (see Methods).

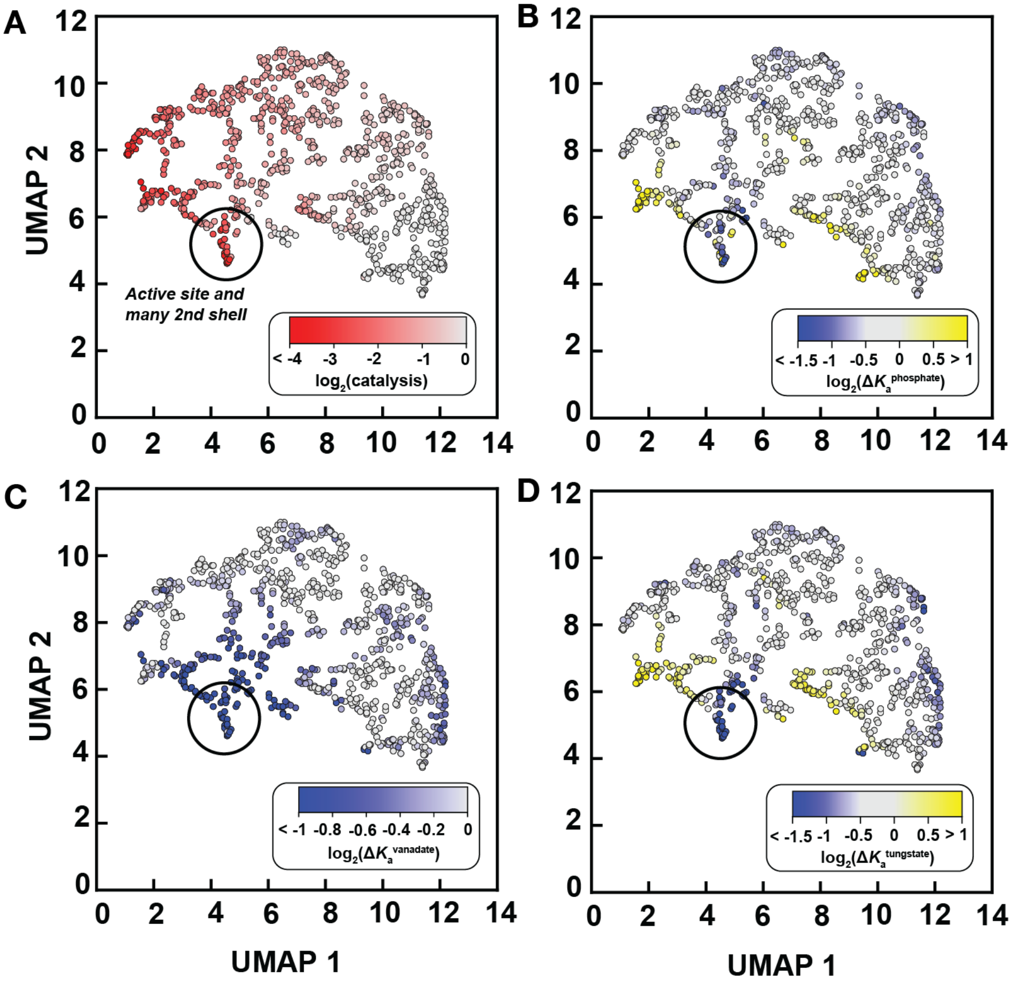


**Fig. S14.** Visualization of relationships within high-dimensional thermodynamic and kinetic data via UMAP dimensionality reduction using log-transformed values for kinetic ($k_{cat}/K_{M,chem}$ for the MeP substrate) and thermodynamic parameters ($K_a$s for phosphate, vanadate, and tungstate) as inputs. UMAPs are colored by the relative impacts of each mutation on: (A) catalysis ($k_{cat}/K_{M,Chem}$), (B) $P_i$ affinity, (C) vanadate affinity, and (D) tungstate affinity. Black circle denotes locations of active site and second shell mutants with catalytic and TSA affinity effects of similar magnitude.

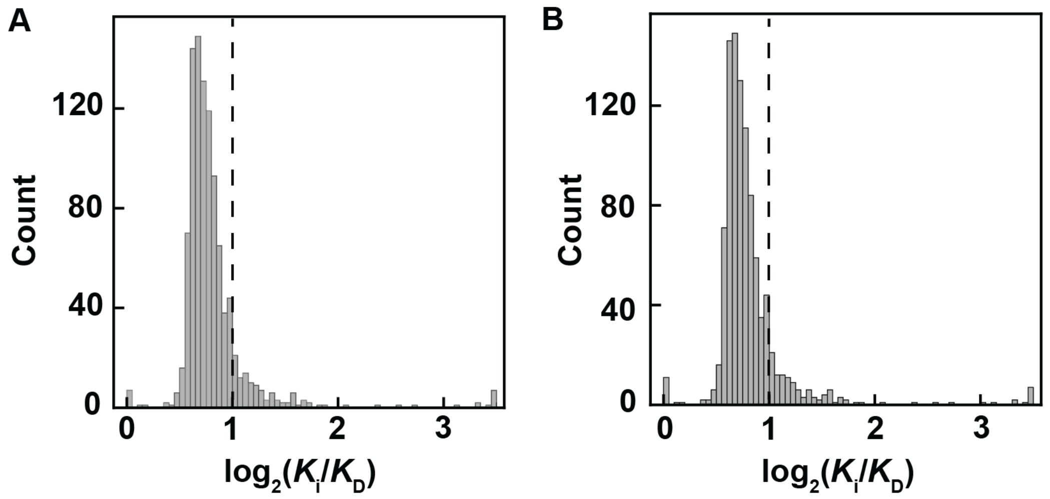


**Fig. S15.** Comparison of median measured $K_i$ values and median $K_D$ values obtained using the Cheng-Prussoff correction (4) for (A) vanadate and (B) tungstate affinity measurements. The dashed line denotes a two-fold change after correction.

**Table S1.** Catalytic and vanadate and tungstate affinity effects (fold change from WT) of active site $Zn^{2+}$ ligand mutants.

| | Catalysis | | Vanadate Affinity | | Tungstate Affinity | |
|---|---|---|---|---|---|---|
| **Mutant** | **Effect** | ***p*-value** | **Effect** | ***p*-value** | **Effect** | ***p*-value** |
| **D38G** | $6.0 \times 10^{1}$ | $2.0 \times 10^{-3}$ | 8.5 | 0.0 | 7.7 | 0.0 |
| **D38V** | – | – | – | – | – | – |
| **D305G** | $1.5 \times 10^{1}$ | $2.2 \times 10^{-2}$ | – | – | $4.1 \times 10^{-2}$ | 0.0 |
| **D305V** | – | – | – | – | – | – |
| **H309G** | 1.3 | $5.0 \times 10^{-1}$ | 4.1 | $8.0 \times 10^{-4}$ | 5.0 | 0.0 |
| **H309V** | $4.7 \times 10^{-1}$ | $7.1 \times 10^{-2}$ | $5.5 \times 10^{1}$ | $1.7 \times 10^{-3}$ | $1.3 \times 10^{1}$ | 0.0 |
| **D352G** | 9.7 | 0.0 | 2.4 | $1.8 \times 10^{-2}$ | 8.2 | 0.0 |
| **D352V** | – | – | – | – | – | – |
| **H353G** | – | – | – | – | – | – |
| **H353V** | 5.0 | $3.1 \times 10^{-1}$ | 1.7 | $2.0 \times 10^{-2}$ | 1.6 | |
| **H486G** | $1.6 \times 10^{1}$ | $7.0 \times 10^{-3}$ | $3.5 \times 10^{1}$ | $5.0 \times 10^{-4}$ | $1.1 \times 10^{1}$ | $1.2 \times 10^{-3}$ |
| **H486V** | 3.1 | $1.7 \times 10^{-1}$ | – | – | 5.8 | $2.9 \times 10^{-3}$ |

**Table S2.** Breakdown of catalytic effects and vanadate affinity effects by interaction shell.

| | Catalytic (down) & Affinity Effect | | Catalytic (up) & Affinity Effect | | Catalytic Effect Only | | Affinity Effect Only | | No Effect | Missing or Limit | Total |
|---|---|---|---|---|---|---|---|---|---|---|---|
| Shell | Tighter | Weaker | Tighter | Weaker | Down | Up | Tighter | Weaker | | | |
| Active Site | 0 | 2 | 0 | 0 | 1 | 0 | 0 | 0 | 0 | 5 | 8 |
| 2 | 0 | 11 | 0 | 0 | 5 | 3 | 1 | 9 | 10 | 11 | 50 |
| 3 | 0 | 12 | 0 | 1 | 14 | 7 | 0 | 10 | 81 | 13 | 138 |
| 4 | 0 | 9 | 0 | 1 | 40 | 2 | 0 | 10 | 196 | 18 | 276 |
| 5 | 0 | 6 | 0 | 0 | 45 | 10 | 0 | 7 | 243 | 15 | 326 |
| 6 | 0 | 1 | 0 | 0 | 21 | 2 | 0 | 3 | 162 | 9 | 198 |
| 7 | 0 | 0 | 0 | 0 | 3 | 0 | 0 | 0 | 30 | 2 | 40 |
| 8 | 0 | 0 | 0 | 0 | 0 | 0 | 0 | 0 | 2 | 0 | 2 |
| 9 | 0 | 0 | 0 | 0 | 0 | 0 | 0 | 0 | 2 | 0 | 2 |
| N.D.[a] | 0 | 0 | 0 | 0 | 0 | 0 | 0 | 0 | 12 | 0 | 12 |

a. N.D. denotes residues for which the interaction shell cannot be determined, as these are residues are unmodeled in the WT PafA crystal structure.

*Data for each mutant are located in Dataset S1, and in the Open Science Foundation data repository (https://osf.io/k8uer/)

**Table S3.** Fractions of catalytic effects and vanadate affinity effects for measurable mutants by interaction shell.

| | Catalytic (down) & Affinity Effect | | Catalytic (up) & Affinity Effect | | Catalytic Effect Only | | Affinity Effect Only | | No Effect |
|---|---|---|---|---|---|---|---|---|---|
| Shell | Tighter | Weaker | Tighter | Weaker | Down | Up | Tighter | Weaker | |
| Active Site | 0 | 0.67 | 0 | 0 | 0.33 | 0 | 0 | 0 | 0 |
| 2 | 0 | 0.28 | 0 | 0 | 0.13 | 0.08 | 0.03 | 0.23 | 0.26 |
| 3 | 0 | 0.10 | 0 | 0.01 | 0.11 | 0.06 | 0 | 0.08 | 0.65 |
| 4 | 0 | 0.04 | 0 | 0 | 0.16 | 0.01 | 0 | 0.04 | 0.77 |
| 5 | 0 | 0.02 | 0 | 0 | 0.14 | 0.03 | 0 | 0.02 | 0.78 |
| 6 | 0 | 0.01 | 0 | 0 | 0.11 | 0.01 | 0 | 0.02 | 0.86 |
| 7 | 0 | 0 | 0 | 0 | 0.21 | 0 | 0 | 0 | 0.79 |
| 8 | 0 | 0 | 0 | 0 | 0 | 0 | 0 | 0 | 0 |
| 9 | 0 | 0 | 0 | 0 | 0 | 0 | 0 | 0 | 0 |
| N.D.[a] | 0 | 0 | 0 | 0 | 0 | 0 | 0 | 0 | 0 |

a. N.D. denotes residues for which the interaction shell cannot be determined, as these are residues are unmodeled in the WT PafA crystal structure.

*Data for each mutant are located in Dataset S1, and in the Open Science Foundation data repository (https://osf.io/k8uer/)

**Table S4**. Breakdown of catalytic effects and tungstate affinity effects by interaction shell.

| | Catalytic (down) & Affinity Effect | | Catalytic (up) & Affinity Effect | | Catalytic Effect Only | | Affinity Effect Only | | No Effect | Missing or Limit | Total |
|---|---|---|---|---|---|---|---|---|---|---|---|
| **Shell** | **Tighter** | **Weaker** | **Tighter** | **Weaker** | **Down** | **Up** | **Tighter** | **Weaker** | | | |
| Active Site | 0 | 2 | 0 | 0 | 2 | 0 | 0 | 0 | 0 | 4 | 8 |
| 2 | 3 | 12 | 0 | 1 | 3 | 2 | 3 | 8 | 9 | 9 | 50 |
| 3 | 6 | 10 | 2 | 2 | 9 | 4 | 16 | 9 | 60 | 20 | 138 |
| 4 | 25 | 5 | 2 | 0 | 19 | 2 | 32 | 9 | 162 | 20 | 276 |
| 5 | 27 | 2 | 1 | 0 | 24 | 9 | 21 | 9 | 218 | 15 | 326 |
| 6 | 3 | 1 | 0 | 0 | 18 | 2 | 12 | 3 | 144 | 15 | 198 |
| 7 | 1 | 0 | 0 | 0 | 6 | 0 | 1 | 3 | 25 | 4 | 40 |
| 8 | 0 | 0 | 0 | 0 | 0 | 0 | 0 | 0 | 2 | 0 | 2 |
| 9 | 0 | 0 | 0 | 0 | 0 | 0 | 0 | 0 | 2 | 0 | 2 |
| N.D.[a] | 0 | 0 | 0 | 0 | 0 | 0 | 0 | 1 | 10 | 1 | 12 |

a. N.D. denotes residues for which the interaction shell cannot be determined, as these are residues are unmodeled in the WT PafA crystal structure.

**Table S5**. Fraction of catalytic effects and tungstate affinity effects by interaction shell.

| | Catalytic (down) & Affinity Effect | | Catalytic (up) & Affinity Effect | | Catalytic Effect Only | | Affinity Effect Only | | No Effect |
|---|---|---|---|---|---|---|---|---|---|
| Shell | Tighter | Weaker | Tighter | Weaker | Down | Up | Tighter | Weaker | |
| Active Site | 0 | 0.5 | 0 | 0 | 0.5 | 0 | 0 | 0 | 0 |
| 2 | 0.07 | 0.29 | 0 | 0.02 | 0.07 | 0.05 | 0.07 | 0.2 | 0.22 |
| 3 | 0.05 | 0.09 | 0.02 | 0.02 | 0.08 | 0.03 | 0.14 | 0.08 | 0.51 |
| 4 | 0.1 | 0.02 | 0.01 | 0 | 0.07 | 0.01 | 0.13 | 0.04 | 0.64 |
| 5 | 0.09 | 0.01 | 0 | 0 | 0.08 | 0.03 | 0.07 | 0.03 | 0.70 |
| 6 | 0.02 | 0.01 | 0 | 0 | 0.1 | 0.01 | 0.07 | 0.02 | 0.79 |
| 7 | 0.03 | 0 | 0 | 0 | 0.17 | 0 | 0.03 | 0.08 | 0.69 |
| 8 | 0 | 0 | 0 | 0 | 0 | 0 | 0 | 0 | 1 |
| 9 | 0 | 0 | 0 | 0 | 0 | 0 | 0 | 0 | 1 |
| N.D.[a] | 0 | 0 | 0 | 0 | 0 | 0 | 0 | 0.09 | 0.91 |

a. N.D. denotes residues for which the interaction shell cannot be determined, as these are residues are unmodeled in the WT PafA crystal structure.

**Table S6.** Breakdown of glycine catalytic effects and tungstate affinity effects by interaction shell.

| | Catalytic (down) & Affinity Effect | | Catalytic (up) & Affinity Effect | | Catalytic Effect Only | | Affinity Effect Only | | No Effect | Missing or Limit | Total |
|---|---|---|---|---|---|---|---|---|---|---|---|
| **Shell** | **Tighter** | **Weaker** | **Tighter** | **Weaker** | **Down** | **Up** | **Tighter** | **Weaker** | | | |
| Active Site | 0 | 1 | 0 | 0 | 1 | 0 | 0 | 0 | 0 | 2 | 4 |
| 2 | 2 | 3 | 0 | 1 | 1 | 2 | 1 | 5 | 5 | 5 | 25 |
| 3 | 2 | 3 | 0 | 1 | 3 | 2 | 9 | 1 | 29 | 19 | 69 |
| 4 | 14 | 3 | 2 | 0 | 1 | 1 | 19 | 0 | 75 | 23 | 138 |
| 5 | 15 | 0 | 0 | 0 | 3 | 5 | 17 | 0 | 101 | 22 | 163 |
| 6 | 3 | 0 | 0 | 0 | 5 | 2 | 9 | 0 | 65 | 15 | 99 |
| 7 | 1 | 0 | 0 | 0 | 0 | 0 | 1 | 0 | 11 | 7 | 20 |
| 8 | 0 | 0 | 0 | 0 | 0 | 0 | 0 | 0 | 1 | 0 | 1 |
| 9 | 0 | 0 | 0 | 0 | 0 | 0 | 0 | 0 | 1 | 0 | 1 |
| N.D.[a] | 0 | 0 | 0 | 0 | 0 | 0 | 0 | 0 | 5 | 0 | 5 |

a. N.D. denotes residues for which the interaction shell cannot be determined, as these are residues are unmodeled in the WT PafA crystal structure.

**Table S7**. Breakdown of valine catalytic effects and tungstate affinity effects by interaction shell.

| | Catalytic (down) & Affinity Effect | | Catalytic (up) & Affinity Effect | | Catalytic Effect Only | | Affinity Effect Only | | No Effect | Missing or Limit | Total |
|---|---|---|---|---|---|---|---|---|---|---|---|
| Shell | Tighter | Weaker | Tighter | Weaker | Down | Up | Tighter | Weaker | | | |
| Active Site | 0 | 1 | 0 | 0 | 1 | 0 | 0 | 0 | 0 | 2 | 4 |
| 2 | 1 | 8 | 0 | 0 | 2 | 0 | 1 | 3 | 3 | 7 | 25 |
| 3 | 3 | 7 | 2 | 1 | 4 | 2 | 6 | 8 | 24 | 12 | 69 |
| 4 | 10 | 2 | 0 | 0 | 14 | 1 | 11 | 9 | 69 | 22 | 138 |
| 5 | 11 | 2 | 1 | 0 | 19 | 4 | 3 | 9 | 98 | 16 | 163 |
| 6 | 0 | 1 | 0 | 0 | 13 | 0 | 1 | 3 | 73 | 8 | 99 |
| 7 | 0 | 0 | 0 | 0 | 5 | 0 | 0 | 3 | 9 | 3 | 20 |
| 8 | 0 | 0 | 0 | 0 | 0 | 0 | 0 | 0 | 1 | 0 | 1 |
| 9 | 0 | 0 | 0 | 0 | 0 | 0 | 0 | 0 | 1 | 0 | 1 |
| N.D.[a] | 0 | 0 | 0 | 0 | 0 | 0 | 0 | 1 | 4 | 0 | 5 |

a. N.D. denotes residues for which the interaction shell cannot be determined, as these are residues are unmodeled in the WT PafA crystal structure.

**Table S8.** Numbers of glycine scanning library mutants with TSA and $P_i$ affinity effects.

| | Catalytic Effect | | | No Catalytic Effect | | | Missing |
|---|---|---|---|---|---|---|---|
| **Inhibitor** | **Weaker** | **Unchanged** | **Stronger** | **Weaker** | **Unchanged** | **Stronger** | |
| Vanadate | 17 | 43 | 0 | 25 | 409 | 1 | 22 |
| Tungstate | 10 | 14 | 38 | 8 | 336 | 63 | 48 |
| Phosphate | 8 | 23 | 32 | 22 | 226 | 190 | 16 |

**Table S9.** Numbers of valine scanning library mutants with TSA and $P_i$ affinity effects.

| | Catalytic Effect | | | No Catalytic Effect | | | Missing |
|---|---|---|---|---|---|---|---|
| **Inhibitor** | **Weaker** | **Unchanged** | **Stronger** | **Weaker** | **Unchanged** | **Stronger** | |
| Vanadate | 24 | 91 | 0 | 15 | 352 | 0 | 37 |
| Tungstate | 22 | 67 | 27 | 36 | 312 | 27 | 28 |
| Phosphate | 22 | 62 | 32 | 21 | 280 | 76 | 26 |

**Movie S1.** Three-dimensional PafA structure (5TJ3, (5)), first showing positions with mutational effects on the chemical step of catalysis (red spheres), followed by a comparison of the effects of these mutations on vanadate affinity. All significant catalytic effects are initially presented (red), followed by the subset of these mutations that have weaker-than-WT vanadate affinities (purple), then the subset that have unchanged inhibitor affinities (red), and finally these sets shown together. Numbers of mutations in each category are given in *SI Appendix*, Table S8, and all data are given in Dataset S1.

**Movie S2.** Three-dimensional PafA structure (5TJ3, (5)), first showing positions with mutational effects on the chemical step of catalysis (red spheres), followed by a comparison of the effects of these same mutations on tungstate affinity. All significant catalytic effects are initially presented (red), followed by the subset of these mutations that have weaker-than-WT tungstate affinities (purple), the subset that have unchanged inhibitor affinities (red), the subset that have stronger-than-WT affinities (yellow), and finally these sets shown together. Numbers of mutations in each category are given in *SI Appendix*, Table S8, and all data are given in Dataset S1.

**Movie S3.** Three-dimensional PafA structure (5TJ3, (5)), first showing positions with mutational effects on the chemical step of catalysis (red spheres), followed by a comparison of the effects of these same mutations on $P_i$ affinity. All significant catalytic effects are initially presented (red), followed by the subset of these mutations that have weaker-than-WT $P_i$ affinities (purple), the subset that have unchanged inhibitor affinities (red), and the subset that have stronger-than-WT affinities (yellow), followed by these sets shown together. Numbers of mutations in each category are given in *SI Appendix*, Table S8, and all data are given in Dataset S1.

**Dataset S1 (separate file).** Median mutational effects on the chemical step of catalysis ($k_{cat}/K_{M,Chem}$) (from (1)), and the median $K_D$ for phosphate (1), vanadate and tungstate, the $log_2$ fold-change of each of these affinities from wild-type, measures of statistical significance (*p*-values) and whether these measurements are limits, and their interaction shell, for wild-type PafA and each mutant.

Columns in the Dataset are labeled as follows:

MeP_kcat/KMAggLimit: effects on the chemical step of catalysis ($k_{cat}/K_{M,Chem}$)

Log2FC_MeP_kcat/KM: $log_2$ fold-change from wild-type of $k_{cat}/K_{M,Chem}$

MeP_kcat/KMChemBootstrapHypothesisTest: statistical significance of effects (*p*-values) on the chemical step of catalysis ($k_{cat}/K_{M,Chem}$)

Catalytic_Fa_Limit: denotes the presence and type of limit of the measurements.

MO4_KdNormLimitValueMedian, where M denotes P, V, or W: median inhibitor $K_D$s for each mutant.

Log2FC_MO4_KdNormLimitValueMedian: $log_2$ fold-changes from wild-type

MO4_KdNormBootstrapHypothesisTest: statistical significance of $K_D$ effects (p-values)

MO4_KdNormAggLimit: denotes the presence and type of limit on the affinity ($K_D$) measurements (0: not a limit; -1: upper limit; 1: lower limit)

FullShellWithZnLigands: interaction shell of each mutated residue
